# Loss of PV interneurons in the BLA contributes to altered network and behavioral states in chronically epileptic mice

**DOI:** 10.1101/2023.12.05.570112

**Authors:** Phillip L.W. Colmers, Pantelis Antonoudiou, Trina Basu, Garrett Scapa, Patrick Fuller, Jamie Maguire

## Abstract

Psychiatric disorders, including anxiety and depression, are highly comorbid in people with epilepsy. However, the mechanisms mediating the shared pathophysiology are currently unknown. There is considerable evidence implicating the basolateral amygdala (BLA) in the network communication of anxiety and fear, a process demonstrated to involve parvalbumin-positive (PV) interneurons. The loss of PV interneurons has been well described in the hippocampus of chronically epileptic mice and in postmortem human tissue of patients with temporal lobe epilepsy (TLE). We hypothesize that a loss of PV interneurons in the BLA may contribute to comorbid mood disorders in epilepsy. To test this hypothesis, we employed a ventral intrahippocampal kainic acid (vIHKA) model of chronic epilepsy in mice, which exhibits profound behavioral deficits associated with chronic epilepsy. We demonstrate a loss of PV interneurons and dysfunction of remaining PV interneurons in the BLA of chronically epileptic mice. Further, we demonstrate altered principal neuron function and impaired coordination of BLA network and behavioral states in chronically epileptic mice. To determine whether these altered network and behavioral states were due to the loss of PV interneurons, we ablated a similar percentage of PV interneurons observed in chronically epileptic mice by stereotaxically injecting AAV-Flex-DTA into the BLA of PV-Cre mice. Loss of PV interneurons in the BLA is sufficient to alter behavioral states, inducing deficits in fear learning and recall of fear memories. These data suggest that compromised inhibition in the BLA in chronically epileptic mice contributes to behavioral deficits, suggesting a novel mechanism contributing to comorbid anxiety and epilepsy.

**Significance Statement:** Psychiatric illnesses and epilepsy are highly comorbid and negatively impact the quality of life of people with epilepsy. The pathophysiological mechanisms mediating the bidirectional relationship between mood disorders and epilepsy remain unknown and, therefore, treatment options remain inadequate. Here we demonstrate a novel mechanism, involving the loss of PV interneurons in the BLA, leading to a corruption of network and behavioral states in mice. These findings pinpoint a critical node and demonstrate a novel cellular and circuit mechanism involved in the comorbidity of psychiatric illnesses and epilepsy.

## Introduction

Psychiatric illnesses are highly comorbid in epilepsy, occurring in up to 75% of people with epilepsy (LaFrance et al., 2008). Depression and anxiety are the most prevalent psychiatric comorbidities in epilepsy, with an incidence of 55% and 25-50%, respectively (Brandt et al., 2010; Lambert and Robertson, 1999). Thus, the incidence is much higher than the general population or in patients with other chronic illnesses (Josephson and Jetté, 2017). Evidence suggests that there is a bidirectional relationship between psychiatric disorders and epilepsy and is thought to involve an unknown shared neurobiological mechanism (Kanner and Balabanov, 2002; Kanner et al., 2018; Mula, 2012). The goal of the current study is to investigate a novel potential mechanism contributing to the comorbidity mediating the bidirectional relationship between mood disorders in epilepsy.

The amygdala plays an integral role in emotional processing (LeDoux, 1992) and has been implicated in the pathophysiology of mood disorders (Price and Drevets, 2009; Ressler, 2010). Computational functions of the amygdala are subserved by oscillations and extensive evidence demonstrates a role for specific oscillatory states in the mPFC and BLA in governing behavioral outcomes (Antonoudiou, 2022; Davis et al., 2017; Felix-Ortiz et al., 2016; Likhtik et al., 2014; Ozawa et al., 2020; Stujenske et al., 2014) (for review see (Tovote et al., 2015)). Oscillations have been shown to be generated by neuronal synchrony that is orchestrated by GABAergic interneurons (Buzsáki and Wang, 2012). In particular, parvalbumin-positive (PV) interneurons have been shown to be critical for the generation of gamma oscillations (Bartos et al., 2007). In the basolateral amygdala, oscillations are driven by GABAergic signaling and PV interneurons play a critical role in generating oscillations in this network (Antonoudiou, 2022). Further, PV interneurons have been shown to be required for the network communication of fear (Davis et al., 2017). Thus, it is well established that PV interneurons in the BLA play a critical role in generating network and behavioral states.

Loss of GABAergic interneurons has been demonstrated in both experimental models of epilepsy as well as in people with epilepsy, particularly in the hippocampus (Liu et al., 2014). A loss of interneurons has also been demonstrated in the amygdala in both in patients with TLE and in experimental epilepsy models (Callahan et al., 1991; Pitkanen et al., 1998; Tuunanen et al., 1996) (nicely reviewed in (Aroniadou-Anderjaska et al., 2008)). In particular, a loss of somatostatin-positive interneurons in the amygdala has been demonstrated in experimental epilepsy models (Sperk et al., 1986; Tuunanen et al., 1997). However, there is a discrepancy between the magnitude of the loss of GABA immunoreactive interneurons and SST interneurons (Pitkanen et al., 1998), suggesting that other subclasses of interneurons may be affected. Although there is extensive evidence for a loss and unique vulnerability of PV interneurons in the hippocampus of human TLE and in experimental models of epilepsy (Bouilleret et al., 2000; Houser, 2014; Marx et al., 2013), surprisingly few studies have investigated the potential loss of PV interneurons in the amygdala. PV interneurons make up nearly 50% of GABAergic interneurons in the rodent BLA (McDonald et al., 2012; Rainnie et al., 2006; Sosulina et al., 2006) (for review see (Capogna, 2014)); whereas, SST interneurons only represent about 5% of this population (Rainnie et al., 2006). Here we demonstrate a loss of PV interneurons in the BLA of chronically epileptic mice, consistent with limited clinical findings suggesting abnormalities in PV interneurons in patients with temporal lobe epilepsy (TLE) (Yilmazer-Hanke et al., 2007). Given the importance of PV interneurons in the BLA for orchestrating network and behavioral states, we proposed that this loss of PV interneurons in the BLA may contribute to comorbid behavioral deficits in chronic epilepsy.

The current study tests the hypothesis that the loss of PV interneurons in the BLA corrupts oscillatory states in the BLA, leading to a breakdown in the network coordination of behavioral states, contributing to an increase in behavioral deficits in chronically epileptic mice. Here we demonstrate that chronically epileptic mice exhibit profound behavioral deficits, consistent with previous reports (Zeidler et al., 2018), including increased avoidance behaviors and anhedonia. Chronically epileptic mice exhibit a substantial reduction in the number of PV interneurons in the BLA and altered inhibitory signaling. The reduction in the number and dysfunction of PV interneurons in the BLA is associated with altered network activity in the BLA. Ablation of an equivalent number of PV interneurons in the BLA to that which is observed in chronically epileptic mice is sufficient to induce behavioral deficits in non-epileptic mice. These data suggest that degeneration of PV interneurons in the BLA of chronically epileptic mice may contribute to comorbid behavioral deficits.

## Materials and Methods

### Animals

Adult male C57BL/6J (Stock #000664), PV-Cre (Stock #017320), and PV-tdTomato (Stock#027395) mice, aged 10-12 weeks old, were purchased from Jackson Lab and group housed (4/cage) in temperature and humidity-controlled housing rooms on a 12-hour light-dark cycle (lights on at 7AM) with ad libitum food and water. Animals were handled according to protocols and procedures approved by the Tufts University Institutional Animal Care and Use Committee (IACUC).

### Behavior paradigms

Open field: Avoidance behaviors in the open field test were measured as previously described (Walton, 2022; Antonoudiou, 2022; Basu, 2022). Briefly, mice were individually placed into the center of the open arena (40 cm x 40 cm) equipped with a photobeam frame with 16 x 16 equally spaced photocells (Hamilton-Kinder). The number of beam breaks, entries, time spent, and distance traveled in the center and the periphery of the apparatus were automatically measured using the Motor Monitor software (Hamilton-Kinder) over the 10 min testing period.

Light/dark box: Avoidance behaviors in the light/dark box were measured as previously described (Melon, 2019; Walton, 2022; Antonoudiou, 2022; Basu, 2022). Briefly, mice were placed individually into the dark compartment of the two-chamber light/dark box apparatus (22 cm x 43 cm) equipped with a photobeam frame with 8 equally spaced photocells (Hamilton-Kinder). Beam breaks, number of entries, time spent, and distance traveled in the light and dark compartments was automatically measured using Motor Monitor software (Hamilton-Kinder) during the 10 min testing period.

Elevated plus maze: The elevated plus maze test was conducted as previously described (Melon, 2018; Antonoudiou, 2022; Walton, 2022). Briefly, mice were individually placed into the center of the elevated plus maze which consists of two opposing 38 cm × 6.5 cm wide arms, standing 75 cm from the ground, one arm exposed and one with closed walls (10 cm high). All arms of the elevated plus maze are equipped with 48 equally spaced photocells allowing automated measurements (MotorMonitor software; Hamilton-Kinder) of beam breaks, number of entries, distance traveled, and total time spent in the open and closed arms during the 10 min test.

Sucrose preference: Anhedonia was measured as previously described by our laboratory (Antonoudiou, 2022; Basu, 2022). Mice were individually housed and given ad libitum access to two water bottles, one filled with water and the other filled with 2% sucrose (w/v), for 7 consecutive days. The position of the water bottles were alternated daily to avoid placement preference. The amount of water and sucrose solution consumed was measured daily and sucrose preference was measured as the percentage of sucrose consumed compared to total volume consumed.

Social interaction: The social interaction test was performed using a three-chamber apparatus (34”Wx11 ½”Dx14”H) in which two wire-mesh cylinders (4”Dx4”H) were placed in each of the left and right chambers. The mice were individually placed into the center of the apparatus and were allowed to habituate and explore the environment for 5 mins. After the 5 min habituation session, an unfamiliar conspecific mouse was placed into one of the cylinders. The position of the mouse was alternated between sessions to avoid a place preference bias. During the test session, mice were placed individually into the center of the apparatus and the amount of time spent in the interaction zone (6”D) of the cylinder containing the conspecific mouse was measured during a 10 min period. A social interaction ratio was calculated as the ratio of time spent in the interaction zone in the target session relative to the empty cylinder.

Fear conditioning: Fear conditioning was performed as previously described (Lee, 2016; Ozawa, 2020; Davis, 2018). Mice were individually placed into the fear conditioning chambers (Coulbourn Instruments; H10-11R-TC, 12”Wx10”Dx12”H) and were subjected to a series of three 20 sec tones (2800 Hz, 80 dB) which ended simultaneously with foot shocks (2s, 0.7 mA) separated by a 1 min interval over a 6 min testing period. Twenty-four hours later, the recall of the contextual fear memory was measured by returning the mice to the fear conditioning chambers for a 3 min period and the amount of freezing behavior was measured in the absence of the tone or shock (contextual recall). Three hours later, cued fear conditioning was measured in a novel environment, a rectangular plastic container with black and white stripes along the sides and bedding scented with 1% acetic acid, in response to presentation with the same tone protocol that was presented during the training phase (3 minute baseline and three 20 s tones separated by a 60 s gap) without shocks (cued recall). Freezing behavior was analyzed using Actimetrics FreezeFrame software (Coulbourn Instruments, bout length 1s). The percent time freezing for 40 secs after each tone was calculated as a measure of cued fear memory. The freezing behavior during the training phase was also measured to assess any potential baseline differences in freezing behavior unrelated to memory.

### Immunohistochemistry

Immunohistochemistry was performed as previously described (Hooper, 2016; Melon, 2019; DiLeo, 2022). Mice were anesthetized with isoflurane, decapitated, and the brain rapidly extracted. The brain was fixed by immersion fixation overnight in 4% PFA at 4°C and then cryoprotected in 10-30% sucrose. The brains were then flash frozen in isopentane on dry ice and stored at -80°C until cryosectioning. Free floating 40 μm coronal slices were incubated in a monoclonal antibody against parvalbumin (PV) (1:1000, Sigma P3088) for 24 hours at 4°C followed by incubation with a biotinylated goat anti-mouse (1:200, ThermoFisher Scientific A28181) antibody for two hours at room temperature and then a streptavidin conjugated Alexa-Fluor 488 (1:200, ThermoFisher Scientific S32354) for two hours at room temperature. Slices were mounted and cover slipped with an antifade hard set mounting medium with DAPI (Vectashield H1500). Fluorescence imaging was performed using a Nikon A1R confocal microscope and the number of PV-positive cells in the basolateral amygdala was measured using Image J software.

### Electrophysiology

Mice were anesthetized with isoflurane, rapidly decapitated, and the brain was rapidly extracted and placed in ice cold slicing solution containing (in mM) 150 sucrose, 15 glucose, 33 NaCl, 25 NaHCO_3_, 2.5 KCl, 1.25 NaH_2_PO_4_, 1 CaCl_2_, 7 MgCl_2_ (300 – 310 mOsm). Coronal sections (350 μm) were prepared on a vibratome and incubated at 33°C in normal artificial cerebral spinal fluid (aCSF) containing (in mM) 126 NaCl, 10 glucose, 2 MgCl_2_, 2 CaCl_2_, 2.5 KCl, 1.25 NaHCO_3_, 1.5 Na-pyruvate, 1 L-glutamine (300 – 310 mOsm) for at least one hour prior to recording. Electrophysiological measurements were performed at 33°C (maintained using in line heater, Warner Instruments) and continuously perfused with aCSF bubbled with 95% O2 and 5% CO2 at a rate of ≥ 4 ml/minute. Voltage clamp and current clamp recordings were performed on visually identified principal neurons in the BLA and fluorescently labeled PV interneurons (PV-tdTomato) using a 200B Axopatch amplifier (Molecular Devices) and recorded using borosilicate glass micropipettes (World Precision Instruments) with DC resistance of 5–8 MΩ and PowerLab Hardware and LabChart 7 data acquisition software (AD Instruments). Spontaneous excitatory postsynaptic currents (sEPSCs) and inh3-ibitory postsynaptic currents (sIPSCs) were recorded in the voltage clamp configuration at -60mV and 0mV, respectively, using a cesium methanesulfonate based internal solution: 140 mM cesium methanesulfonate, 10 mM HEPES, 5 mM NaCl, 0.2 mM EGTA, 2 mM Mg-ATP, and 0.3 mM Na-GTP (pH 7.25; 280–290 mOsm). Synaptic events over a 120 s epoch were analyzed using a custom MATLAB script. The frequency, amplitude, and decay of synaptic events was measured for each cell and averaged across experimental groups. Current clamp recordings were performed in the I=0 mode using a potassium gluconate based internal solution: 130 K-gluconate, 10 KCl, 4 NaCl, 10 HEPES, 0.1 EGTA, 2 Mg-ATP, 0.3 Na-GTP (pH 7.25, 280–290 mOsm/L H_2_O). Input-output curves were generated in response to a series of current injections from 0-150pA in 10pA steps. Rheobase was measured in response to a -70-0mV current ramp. The resonant frequency profiles of principal neurons and PV interneurons were measured in response to a chirp stimulus. Subthreshold membrane properties and threshold firing of each neuron was measured in response to a sinusoidal chirp current of varying frequency from 0-60Hz over a 60s period for the subthreshold chirp and over a 3s or 20s period for the suprathreshold chirp. The maximum amplitude response, peak power, and firing threshold over this frequency range was determined as a measure of the preferred resonant frequency of each neuron and averaged across neurons between experimental groups. Intrinsic electrophysiological properties, including input resistance, impedance, and whole cell time constant were measured for each neuron. Cells were eliminated from analysis if the series resistance was greater than 20 MΩ or changed >20% over the course of the experiment.

Ex vivo LFP recordings from acute brain slices were conducted as previously described (Antonoudiou et al. 2020). Briefly, 350 μm coronal slices collected as above, hemisected with the hippocampus removed, and recovered in interface conditions for one hour before being transferred to an interface recording chamber. The aCSF used for patch clamp recordings was modified for acute LFP recordings to induce gamma oscillations, and contained elevated potassium (7.5 mM KCl) and 800 nM kainic acid. The LFP was recorded through an aCSF-containing borosilicate pipette inserted into the BLA, data were acquired through LabChart (ADInstruments) at 10 KHz, the power line noise (59-61 Hz) was removed with values filled using nearest interpolation, and spectral analysis was performed as below (refer to LFP recordings section).

### Stereotaxic Surgery

Mice were anesthetized with a ketamine (90 mg/kg, i.p.)/xylazine (5-10 mg/kg, i.p.) cocktail and treated with sustained release buprenorphine (0.5-1.0 mg/kg, s.c.) prior to surgical procedures. To generate chronically epileptic mice, 100 nl of 20 mM kainic acid (Sigma-Aldrich, #K0250; dissolved in saline) was stereotaxically injected into the ventral hippocampus (AP -3.60 mm, ML -2.80 mm, DV -2.80 mm from Dura; Zeidler, et al., 2018). Control mice were injected with 100 nL of saline in the ventral hippocampus. For local field potential recordings (LFPs), mice were implanted with a custom head mount (Pinnacle #8201) modified to include a depth electrode (PFA-coated stainless-steel wire, A-M systems) which was stereotaxically implanted into the BLA (AP -1.35 mm, ML 3.30 mm, DV – 4.50 mm from Dura). Stainless steel screws served as a reference lead and an animal ground. EMG wires were also positioned in the neck muscles. Mice were allowed to recover for a minimum of 5 days prior to experimentation. For the PV ablation experiments, mice were stereotaxically injected with either a control virus rAAV2-mCherry or CMV-β-globin-DIO-mCherry-DTA-hGH pA (AAV-Flex-DTA; generated by Dr. Patrick M. Fuller, Harvard Medical School) into the BLA using a Hamilton syringe and the stereotaxic coordinates above. Experiments were performed at 3 weeks following injection to allow for optimal virus expression and PV ablation.

### LFP Recordings

LFP recordings were performed as previously described (Antonoudiou, 2022; DiLeo, 2022). LFP recordings in the BLA and frontal cortex were recorded at 4 KHz and amplified 100X and acquired using Lab Chart software (AD Instruments). For the analysis, the raw data was band pass filtered (1-300 Hz) and spectral analysis was performed using custom analysis script developed in MATLAB using MatWAND (https://github.com/pantelisantonoudiou/MatWAND) (Antonoudiou et al., 2022). Briefly, the recordings were divided into 5 second overlapping segments and the power spectral density for a range of frequencies was obtained (Oppenheim et al., 1999) utilizing a fast Fourier transform similar to previous reports (Kruse and Eckhorn, 1996; Frigo and Johnson, 1998; Pape et al., 1998; Freeman et al., 2000).

For a subset of surgically implanted LFP mice, a brief restraint stress consisting of immobilization in a 50 mL falcon tube with a ¼ inch breathing hole drilled into the end for 30 minutes was performed. LFP recordings were conducted for 1 hour prior to the restraint stress, and mice were returned to their recording chambers following the restraint stress where post-stress LFP recordings were conducted for an additional 2 hours. These post-stress LFP were split into 30 minute segments and normalized to the 1 hour baseline to examine any stress-induced changes in local network function.

### Statistical Analysis

Data were analyzed using Prism 8/10 software (GraphPad) and custom scripts developed in MATLAB (Mathworks). Statistical significance between two experimental groups was determined using a Student’s t-test. For the power analysis across specific frequency bands, a post-hoc Šídák’s multiple comparisons test was performed to determine statistical significance. P values < than 0.05 were considered statistically significant, with 1-4 symbols used in figures to denote a significance level of p<0.05, p<0.01, p<0.001, and p<0.0001 respectively.

## Results

### Behavioral comorbidities in chronically epileptic mice

Chronically epileptic mice were generated following a single 100 nl injection of 20 mM kainic acid unilaterally into the hippocampus of male C57Bl/6J mice. Avoidance behaviors were assessed in chronically epileptic mice 60 days post-kainic acid injection using the open field test, light-dark box, and elevated plus maze. Anhedonia was also assessed using the sucrose preference test. Chronically epileptic male mice exhibit an increase in the amount of time spent in the periphery (556.30±10.57 s) and a decrease in the amount of time spent in the center of the open field (43.66±10.57 s) compared to saline-injected controls (periphery: 500.20±8.18 s; center: 99.78±8.18 s) (Figure 1A; Position x Treatment Interaction: F_(1,19)_=17.10, p=0.0006; 2-way ANOVA; N= 10-11 mice per experimental group). Similarly, chronically epileptic male mice spend an increased amount of time in the dark chamber (539.40±21.86 s) and a decreased amount of time spent in the light chamber of the light-dark box (47.75±18.56 s) compared to controls (dark: 310.4±20.35 s; light: 249.0±19.45 s) (Figure 1B; Position x Treatment Interaction: F_(1,19)_=57.99, p<0.0001, 2-way ANOVA; N=10-11 mice per experimental group). In the elevated plus maze, chronically epileptic male mice make a decreased number of entries into the open arm compared to controls (IHSa: 45.60±9.26 entries; IHKA: 21.00±3.67 entries; p=0.0193; unpaired 2-tail t-test) without a significant difference in the amount of time spent in the open arm (IHSa: 84.17±15.43 s; IHKA: 125.30±24.96 s; p=0.188; unpaired 2-tail t-test) (Figure 1C; N=10-11 mice per experimental group). Collectively, these data demonstrate an increase in avoidance behaviors in chronically epileptic male mice compared to controls. Chronically epileptic mice also exhibit increased anhedonia, exhibited by a decrease in sucrose preference compared to controls (Figure 1D; IHSa: 85.14±3.03%; IHKA: 55.73±5.81%; p=0.0003; unpaired 2-tail t-test; N=10-11 mice per experimental group), with chronically epileptic mice demonstrating no preference relative to chance (50%).

**Figure 1.**
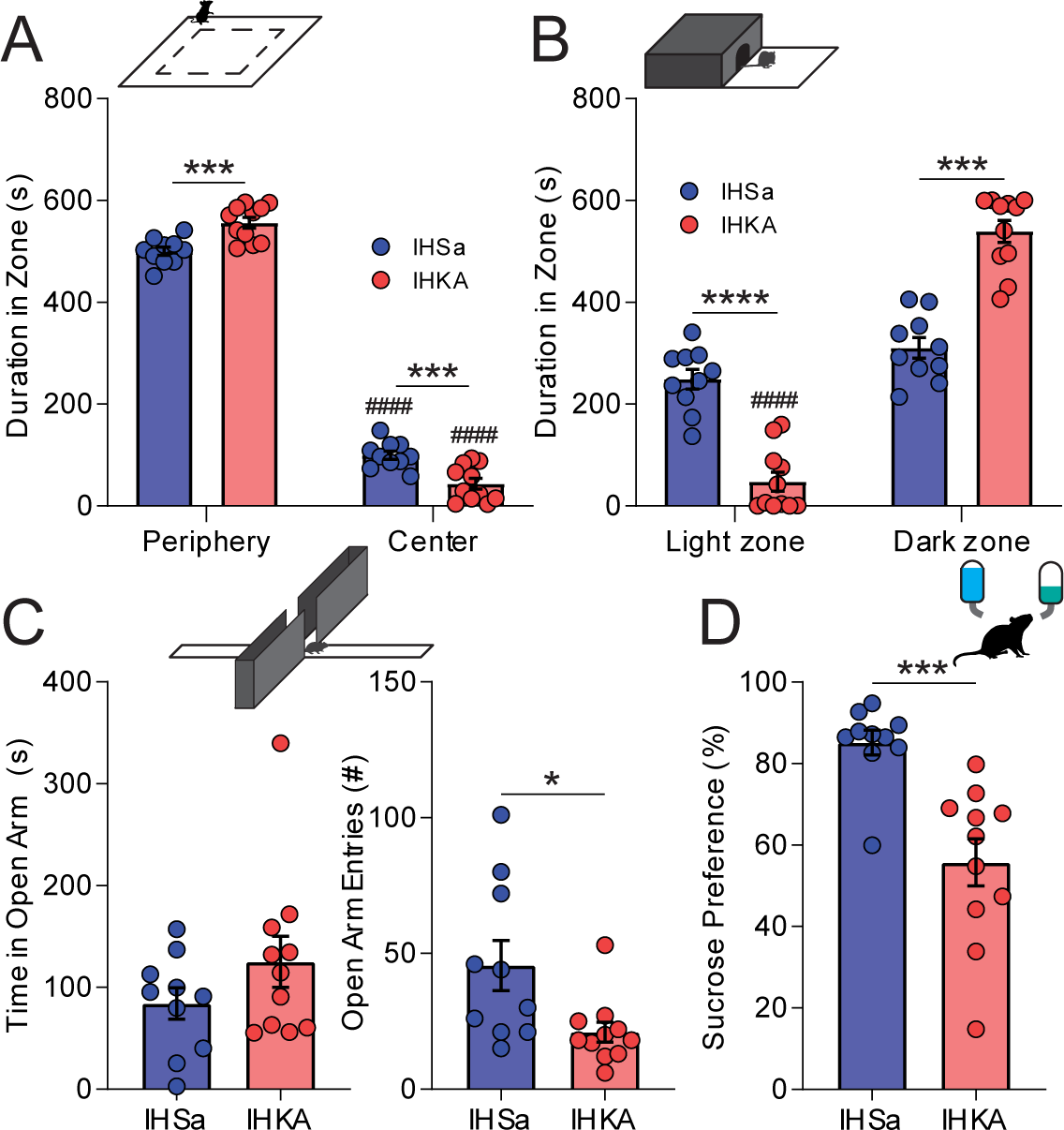
Anxiety-like behaviors in chronically epileptic mice. A, Schematic of the open field test (top) in which both IHSa and IHKA mice spent significantly more time in the periphery of the open field, with IHKA mice spending significantly less time in the center than IHSa mice. B, Schematic of the light dark box (top) in which IHKA mice spent significantly less time in the light zone than IHSa, while IHSa mice spent an equal amount of time in the light and dark zones. C, Schematic of the elevated plus maze (top) in which IHSa mice spent the same amount of time in the open arm as IHKA mice (left), but entered the open arm to explore more frequently (right). D, Schematic of the sucrose preference test (top) showing IHKA mice lack hedonic sucrose-seeking behavior. * denotes the degree of significance between conditions, # denotes the degree of significance within a condition group. A,B derive significance from 2-way ANOVA with Šídák’s multiple comparisons test, C, D derive significance from unpaired 2-tail t-test. Data shown as Mean±SEM.

### PV interneuron deficits in the BLA of chronically epileptic mice

To assess whether there is a loss of PV-positive interneurons in the BLA of chronically epileptic mice, we performed immunohistochemistry for PV in C57Bl/6J males 60 days following intrahippocampal injection of either saline or kainic acid (Figure 2A). The average number of PV interneurons in the BLA is significantly reduced in chronically epileptic mice compared to controls (IHSa: 21.52±0.95 cells/section; IHKA: 13.60±0.73 cells/section; p<0.0001; unpaired 2-tail t-test) (Figure 2B; n=120-122 sections, from N=8 mice per experimental group), which can be appreciated in the representative images from saline-(Figure 2C) and kainic acid-injected mice (Figure 2D).

**Figure 2.**
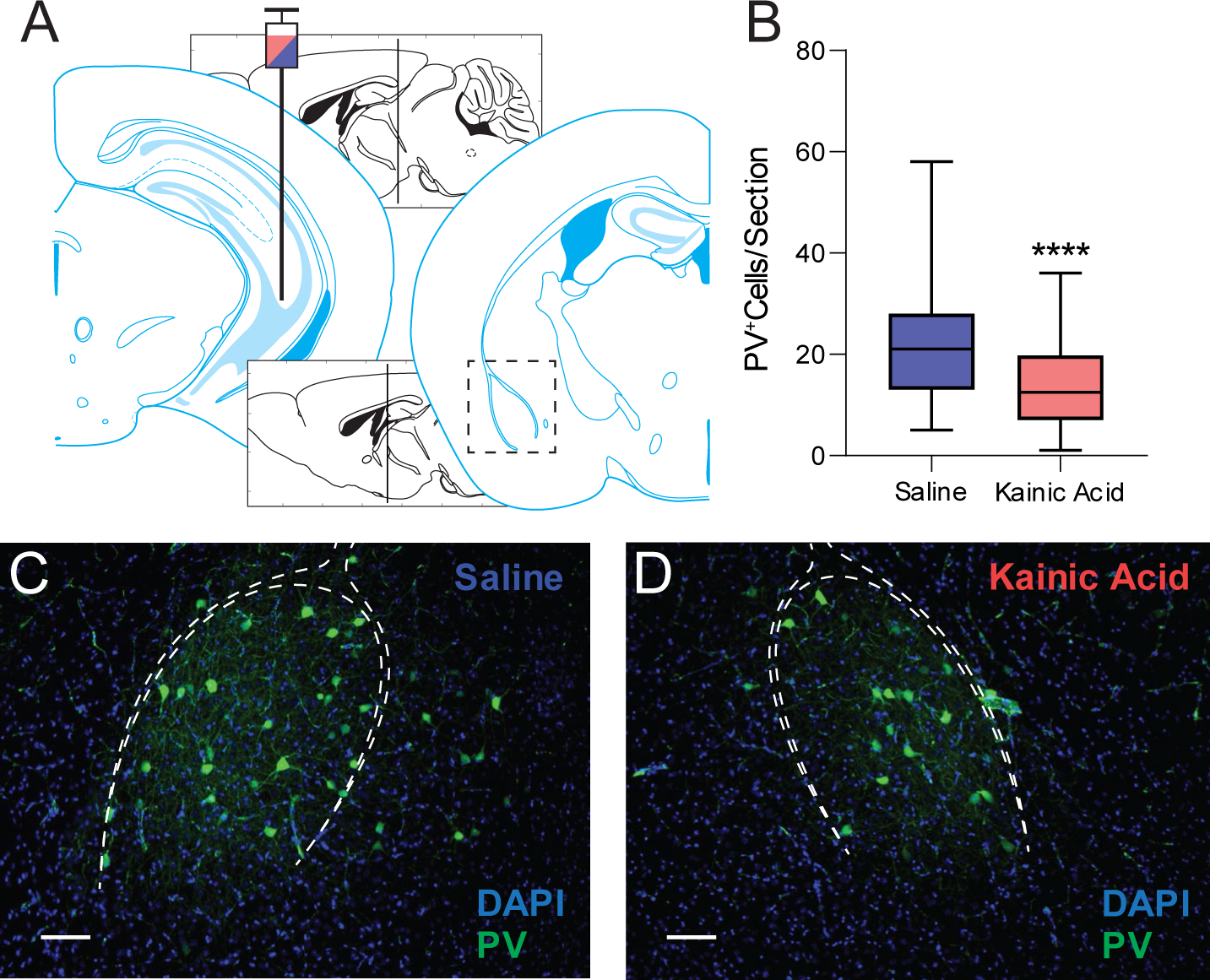
Loss of PV^+^ BLA interneurons following IHKA treatment. A, Schematic showing the intrahippocampal stereotaxic injection site and a slice through the BLA Adapted from Franklin & Paxinos (2008). B, Kainic acid significantly reduced the number of PV^+^ cells counted within the BLA. C, D, Representative fluorescent IHC sections showing a decrease in the number of PV^+^ neurons in the BLA (white dashed outline) of IHKA mice compared to IHSa mice. * denotes the degree of significance between conditions measured by unpaired 2-tail t-test. Data shown as a box-and-whiskers plot with whiskers showing min and max.

The function of remaining PV interneurons in the BLA was assessed using whole cell patch clamp recording. Although there was no difference in the average frequency (Figure 3A inset; IHSa: 16.11±2.42 Hz; IHKA: 15.13±2.47 Hz; p=0.7759; unpaired 2-tail t-test) of spontaneous excitatory postsynaptic currents (sEPSCs) on PV interneurons in the BLA, there was an increase in the average amplitude (Figure 3A insets; IHSa: -31.04±2.39 pA; IHKA: - 40.71±4.15 pA; p=0.0462; unpaired 2-tail t-test; n=31-32 cells; 11-13 mice per experimental group), and an increase in the cumulative distribution of the amplitude and a decrease in the interevent interval (IEI) of sEPSCs (Figure 3A; sEPSC cumulative amplitude: p<0.0001; sEPSC cumulative IEI: p<0.0001; Kolmogorov-Smirnov test). In contrast, there is a significant decrease in the frequency of spontaneous inhibitory postsynaptic current (sIPSCs) in PV interneurons in the BLA of chronically epileptic mice (Figure 3B inset; IHSa: 8.00±1.18 Hz; IHKA: 5.12±0.67 Hz; p=0.0395; unpaired 2-tail t-test) with no change in amplitude (IHSa: 28.47±1.77 pA; IHKA: 28.63±1.23 pA; p=0.9391; unpaired 2-tail t-test) (Figure 3B insets; n=31-32 cells; 11-13 mice per experimental group). In line with this reduction in inhibitory signaling, the cumulative distribution demonstrates a no change in the amplitude and an increase in the IEI of sIPSCs onto PV interneurons in the BLA of chronically epileptic mice (Figure 3B; amplitude: p=0.151; IEI: p<0.0001; Kolmogorov-Smirnov test). However, there is no change in the distribution of the rise or decay times of either sEPSCs or sIPSCs between groups (Figure 3C, D; sEPSC rise: W=-32; p=0.2334; sEPSC decay: W=1; p>0.9999; sIPSC rise: W=-1; p=0.9999; sIPSC decay: W=-20; p=0.7285; Wilcoxon matched pairs signed rank test, 2-tailed).

**Figure 3.**
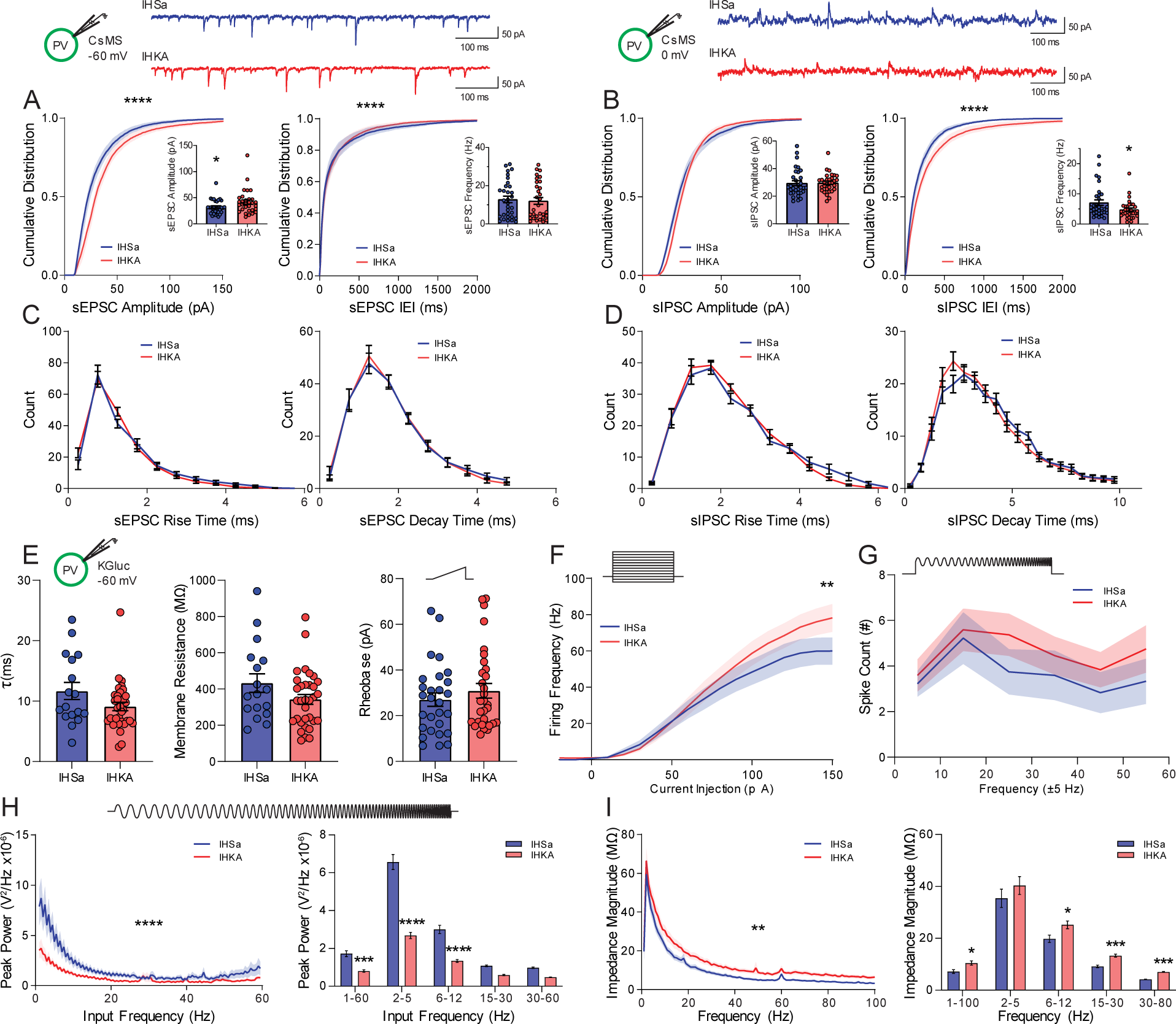
Active and passive electrophysiological properties of genetically identified PV^+^ BLA interneurons. Schematics showing holding current, internal solution and PV-GFP^+^ neuron and representative IHSa and IHKA traces of afferent spontaneous excitatory (A) and inhibitory (B) postsynaptic currents from a whole-cell voltage-clamp recording (top). Excitatory afferent synaptic events (A) in IHKA exhibited a right-shift in the distribution of amplitudes (left) and an increase in the mean amplitude (inset), and a steepening of the distribution of inter-event intervals (IEI, right) with no significant change to the mean frequency of the events (inset) or the rise and decay kinetics (C). Inhibitory afferent synaptic events (B) in IHKA showed no change in the distribution of amplitudes (left) or mean amplitude (inset) but showed a right-shift in the distribution of IEI (right) and a decrease in the mean sIPSC frequency (inset), with no changes to rise or decay kinetics (D). E, Intrinsic membrane properties including the membrane time constant (left), membrane resistance (middle), and rheobase (right) are not significantly altered in IHKA mice. F, In response to successive depolarizing current steps (top), the firing frequency is increased in PV^+^ cells from IHKA mice at higher levels of current injection. G, A Chirp stimulation consisting of an accelerating sine wave on top of a depolarizing current step (top) did not reveal any significant differences between IHSa and IHKA in resonant firing output across the input frequencies tested. H, A subthreshold Chirp current (top) was injected into the current-clamped neuron to determine intrinsic passive resonant membrane properties at different input frequencies, IHKA neurons exhibited reduced resonant properties across the wavelengths tested (left) and within specific low-frequency bands (right). I, Membrane impedance as a function of frequency was significantly higher in IHKA (left) especially in higher frequency bands (right). * denotes the degree of significance between conditions. Cumulative distributions in A, B derive significance from Kolmogorov-Smirnov tests. The mean comparisons in A, B, and E derive significance from unpaired 2-tailed t-tests. Histograms in C, D derive significance from Wilcoxon matched pairs signed rank test, 2-tailed. The two-factor comparisons in F-I derive significance from 2-way ANOVA with Šídák’s multiple comparisons test. Data shown as Mean±SEM.

To evaluate potential changes in the intrinsic properties of PV interneurons, we measured the membrane time constant (Figure 3E, left; IHSa: 11.67±1.41 ms; IHKA: 9.11±0.69 ms; p=0.0716; unpaired 2-tail t-test), membrane resistance (Figure 3E, middle; IHSa: 432.6±51.4 MΩ; IHKA: 342.6±27.4 MΩ; p=0.0955; unpaired 2-tail t-test), and rheobase (Figure 3E, right; IHSa: 27.01±2.94 pA; IHKA: 30.91±3.21 pA; p=0.377; unpaired 2-tail t-test). None of these measures were significantly altered in PV interneurons in BLA slices from control versus chronically epileptic mice (Figure 3E; n=17-33 cells; 5-10 mice per experimental group). Input-output curves in PV interneurons in the BLA demonstrate an increase in the frequency of spikes in response to higher current injections (Figure 3F; Treatment effects: F_(1,913)_=9.39, p=0.0022; 2-way ANOVA; n=29-33 cells, 8-10 mice per experimental group) with a trend towards an increased number of spikes generated in response to the chirp stimulus (accelerating sine wave 0-60 Hz paired with a current step equal to rheobase to initiate firing) in chronically epileptic mice compared to controls (Figure 3G; Treatment effects: F_(1,276)_=2.88, p=0.0906; 2-way ANOVA; n=18-30 cells; 5-10 mice per experimental group).

To assess potential changes in the resonant frequency profiles of PV interneurons to assess their ability to respond to oscillatory inputs, we measured the membrane response to a chirp stimulus. PV interneurons in the BLA slices from chronically epileptic mice exhibit a decrease in peak power across frequencies (1-60 Hz: 8.05×10^-7^±6.30×10^-8^ V^2^/Hz; 2-5 Hz: 2.68×10^-6^±1.66×10^-7^ V^2^/Hz; 6-12 Hz: 1.34×10^-6^±7.89×10^-8^ V^2^/Hz; 15-30 Hz: 5.90×10^-7^±3.04×10^-8^ V^2^/Hz; 30-60 Hz: 4.74×10^-7^±1.68×10^-8^ V^2^/Hz) compared to controls (1-60 Hz: 1.72×10^-6^±1.55×10^-7^ V^2^/Hz; 2-5 Hz: 6.57×10^-6^±3.99×10^-7^ V^2^/Hz; 6-12 Hz: 3.00×10^-6^±2.21×10^-7^ V^2^/Hz; 15-30 Hz: 1.08×10^-6^±4.22×10^-8^ V^2^/Hz; 30-60 Hz: 9.78×10^-7^±4.07×10^-8^ V^2^/Hz) (Figure 3H; Treatment effects: F_(1,290)_=232.3, p<0.0001; 2-way ANOVA; n=27-33 cells, 7-10 mice per experimental group). The impedance across frequencies is increased in PV interneurons in the BLA from chronically epileptic mice compared to controls (Figure 3I; Treatment effects: F_(1,290)_=13.52, p=0.0003; 2-way ANOVA; n=27-33 cells, 7-10 mice per experimental group) with statistical differences measured at 1-100 Hz (IHSa: 9.60±1.00 MΩ; IHKA: 13.92±1.10 MΩ), 6-12 Hz (IHSa: 26.40±1.88 MΩ; IHKA: 33.56±1.98 MΩ), 15-30 Hz (IHSa: 12.19±0.66 MΩ; IHKA: 17.72±0.75 MΩ), and 30-80 Hz (IHSa: 5.55±0.18 MΩ; IHKA: 9.36±0.21 MΩ) (Figure 3I; n=27-33 cells, 7-10 mice per experimental group). These data suggest that the ability of PV interneurons to appropriately respond to oscillatory states or be recruited into generating oscillatory states may be deficient in chronically epileptic mice.

The functional consequences of the loss of and dysfunction of remaining PV interneurons in the BLA was assessed using whole cell patch clamp recording in principal neurons in the BLA. The cumulative distribution of the amplitude of sEPSCs is shifted towards the right and the average amplitude of sEPSCs is increased in principal neurons in the BLA in slices from chronically epileptic mice compared to controls (Supplemental Figure 1A; sEPSC amplitude Distribution: p<0.0001; Kolmogorov-Smirnov test; sEPSC mean amplitudes: IHSa: -19.85±1.50 pA; IHKA: -25.78±1.35 pA; p=0.0073; unpaired 2-tail t-test; n=14-23 cells; 4-5 mice per experimental group) with no change in the distribution of the rise or decay times (Supplemental Figure 1C; sEPSC rise: W=-18; p=0.5186; sEPSC decay: W=13, p=0.5566; Wilcoxon matched pairs rank signed test, 2-tailed; n=14-23 cells, 4-5 mice per experimental group). Although there is no difference in the average frequency of sEPSCs in principal neurons in the BLA of chronically epileptic mice compared to controls, there is a leftward shift in the cumulative distribution of the IEI (Supplemental Figure 1A; sEPSC IEI Distribution: p<0.0001; Kolmogorov-Smirnov test; sEPSC mean frequency: IHSa: 4.08±0.92 Hz; IHKA: 6.56±1.26 Hz; p=0.1701; unpaired 2-tail t-test; n=14-23 cells, 4-5 mice per experimental group).

In contrast, while there is no significant difference in the average amplitude (IHSa: 27.50±1.92 pA; IHKA: 33.62±2.43 pA; p=0.0780; unpaired 2-tail t-test) or frequency (IHSa: 6.90±1.40 Hz; IHKA: 6.96±0.78 Hz; p=0.9680; unpaired 2-tail t-test) of sIPSCs (Supplemental Figure 1B insets; n=14-21 cells, 4-5 mice per experimental group), there is a rightward shift in the cumulative distribution of the peak amplitude of sIPSCs and no change in the IEI in principal BLA neurons in slices from chronically epileptic mice compared to controls (Supplemental Figure 1B; sIPSC cumulative amplitude: p<0.0001; sIPSC cumulative IEI: p=0.1438; Kolmogorov-Smirnov test) with no change in the distribution of rise or decay times (Supplemental Figure 1D; sIPSC rise: W=-16; p=0.5693; sIPSC decay: W=13, p=0.8194; Wilcoxon matched pairs rank signed test, 2-tailed; n=14-21 cells, 4-5 mice per experimental group).

The intrinsic properties of BLA principal neurons are not altered between control and chronically epileptic mice (Supplemental Figure 1E-I). There is no difference in the membrane time constant (IHSa: 13.12±1.15 ms; IHKA: 14.77±1.66 ms; p=0.4330; unpaired 2-tail t-test), membrane resistance (IHSa: 291.3±35.7 MΩ; IHKA: 390.5±43.8 MΩ; p=0.0942; unpaired 2-tail t-test), or rheobase (IHSa: 40.84±4.49 pA; IHKA: 32.20±3.44 pA; p=0.1390; unpaired 2-tail t-test) between control and chronically epileptic mice (Supplemental Figure 1E; n=17-21 cells, 5-7 mice per experimental group). There was also no difference in the input-output curves (Supplemental Figure 1F; Treatment effects: F_(1,612)_=1.90, p=0.169; 2way ANOVA) or number of spikes generated in response to the chirp stimulus (Supplemental Figure 1G; Treatment effects: F_(1,210)_=0.956, p=0.3290; 2way ANOVA; n=18-19 cells, 5-7 mice per experimental group). BLA principal neurons from chronically epileptic mice exhibited enhanced responsivity to the chirp stimulus (1-60 Hz: 6.63×10^-7^±4.24×10^-8^ V^2^/Hz; 2-5 Hz: 1.90×10^-6^±9.35×10^-8^ V^2^/Hz; 6-12 Hz: 1.08×10^-6^±5.39×10^-8^ V^2^/Hz; 15-30 Hz: 5.45×10^-7^±1.49×10^-8^ V^2^/Hz; 30-60 Hz: 4.17×10^-7^±9.05×10^-9^ V^2^/Hz) compared to controls (1-60 Hz: 4.31×10^-7^±4.24×10^-8^ V^2^/Hz; 2-5 Hz: 1.72×10^-6^±1.27×10^-7^ V^2^/Hz; 6-12 Hz: 8.04×10^-7^±4.90×10^-8^ V^2^/Hz; 15-30 Hz: 3.16×10^-7^±1.38×10^-8^ V^2^/Hz; 30-60 Hz: 1.88×10^-7^±2.92×10^-9^ V^2^/Hz) (Supplemental Figure 1H; Treatment effects: F_(1,180)_=38.57, p<0.0001; 2-way ANOVA; n=18-20 cells, 5-7 mice per experimental group) with an increase in impedance observed at 15-30 Hz (IHSa: 7.16±0.45 MΩ; IHKA: 9.13±0.54 MΩ) and 30-80 Hz (IHSa: 3.20±0.11 MΩ; IHKA: 3.92±0.15 MΩ) (Supplemental Figure 1I; Treatment effects F_(1,180)_=1.29, p=0.258; n=18-20 cells, 5-7 mice per experimental group). These data suggest that BLA principal neurons from chronically epileptic mice receive altered synaptic inputs and inappropriately respond to oscillatory inputs.

### Altered BLA network activity in chronically epileptic mice

To examine whether there are changes in BLA network states in chronically epileptic mice, electroencephalogram (EEG) and local field potential recordings (LFPs) were performed in the cortex and BLA, respectively, for four weeks following either IHSa or IHKA injection. The average number of seizures per day (Figure 4A; Wk 1: 0.79±0.38; Wk2: 2.30±0.59; Wk 4: 1.22±0.36; total: 1.39±0.20), seizure duration (Figure 4B; Wk 1: 51.61±1.45 s; Wk 2: 49.11±2.17 s; Wk 4: 56.91±3.23 s; total: 52.60±2.08 s), and total seizure burden (Figure 4C; Wk 1: 105.94±65.95 s; Wk 2: 653.12±151.04 s; Wk 4: 381.56±114.90 s; total: 1140.63±161.11 s) was measured at 1, 2, and 4 weeks post-IHKA injection (N=16 mice). The power of the local field potential across frequencies was compared 1 week (Figure 4D), 2 weeks (Figure 4E), and 4 weeks (Figure 4F) post-IHSa or IHKA injection (N=14-18 mice per experimental group). The changes in the LFP with epilepsy progression can be appreciated from the Power of the LFP subtracted from the LFP at Week 1 (Figure 4G, H, J, K). There are minimal changes in the LFP in the BLA over time in the IHSa group, with only modest increases during week 2 in the 15-30 Hz (Wk 2: 117.3±5.3%; Wk 4: 111.0±13.7%) and 40-70 Hz range (Wk 2: 131.3±8.0%; Wk 4: 108.5±12.8%) (Figure 4I; Week effect: F_(1.34, 81.99)_=3.85, p=0.041; Mixed-effects Analysis; N=12-14 mice). More profound changes in the LFP in the BLA is observed over time with epilepsy progression in IHKA group, with significance in the 6-12 Hz (Wk 2: 122.1±7.7%; Wk 4: 158.5±21.9%), 15-30 Hz (Wk 2: 133.0±10.1%; Wk 4: 170.8±22.3%), 40-70 Hz range (Wk 2: 146.7±11.6%; Wk 4: 196.9±27.7%), and 80-120 Hz (Wk 2: 143.7±13.7%; Wk 4: 156.0±20.6%) (Figure 4L; Week effect: F_(1.351, 114.8)_=37.14, p<0.0001; Mixed-effects Analysis; N=16-20 mice). A direct comparison to changes in individual power bands between IHSa and IHKA mice normalized to their first week (Figure 4N, O, P) revealed changes in epilepsy progression in IHKA mice. There were no significant changes in the 2-5 Hz power band across weeks in both IHSa (Wk 2: 101.85±6.59%; Wk 4: 105.46±9.90%) and IHKA mice (Wk 2: 113.71±9.69%; Wk 4: 122.10±14.70%) (Figure 4N; Treatment effect: F_(1,32)_=1.45, p=0.237; Mixed-effects analysis; N=10-20 mice). Similarly, the IHSa 6-12 Hz Power area (Wk 2: 103.11±5.28%; Wk 4: 99.56±8.77%) was not significantly different in IHKA (Wk 2: 122.08±7.66%; Wk 4: 129.04±16.33%) (Figure 4O; Treatment effect: F_(1,32)_=4.63, p=0.039; Mixed-effects analysis; N=11-20 mice). However, the ratiometric comparison of Power in the 2-5 and 6-12 Hz bands, a marker of BLA network state associated with fear and anxiety, was significantly elevated IHKA (Wk 1: 2.70±0.19; Wk 2: 2.35±0.17; Wk 4: 2.42±0.21) compared to IHSa (Wk 1: 1.43±0.14; Wk 2: 1.73±0.22; Wk 4: 1.60±0.14) (Figure 4M; Week x Treatment Interaction: F_(2,58)_=3.80, p=0.028; Mixed-effects analysis; N=12-20 mice). Power was significantly enhanced at 2 and 4 weeks in the 15-30 Hz band in IHKA mice (Wk 2: 140.17±11.98%; Wk 4: 147.90±17.89%) compared to IHSa mice (Wk 2: 112.97±3.28%; Wk 4: 106.69±4.45%) (Figure 4P; Treatment Effect: F_(1,32)_=5.89, p=0.021; Mixed-effects analysis; N=9-20 mice). Similarly, there were significant differences in power at both 2 and 4 weeks in the 40-70 Hz band in IHKA mice (Wk 2: 175.90±19.02%; Wk 4: 196.93±27.68%) compared to IHSa (Wk 2: 131.33±8.04%; Wk 4: 117.42±10.02%) (Figure 4Q; Week x Treatment Interaction: F_(2,57)_=4.34, p=0.018; Mixed-effects analysis; N=12-20 mice). These data suggest BLA network dysfunction associated with epilepsy progression, which likely involves the loss of PV-positive interneurons in the BLA.

**Figure 4.**
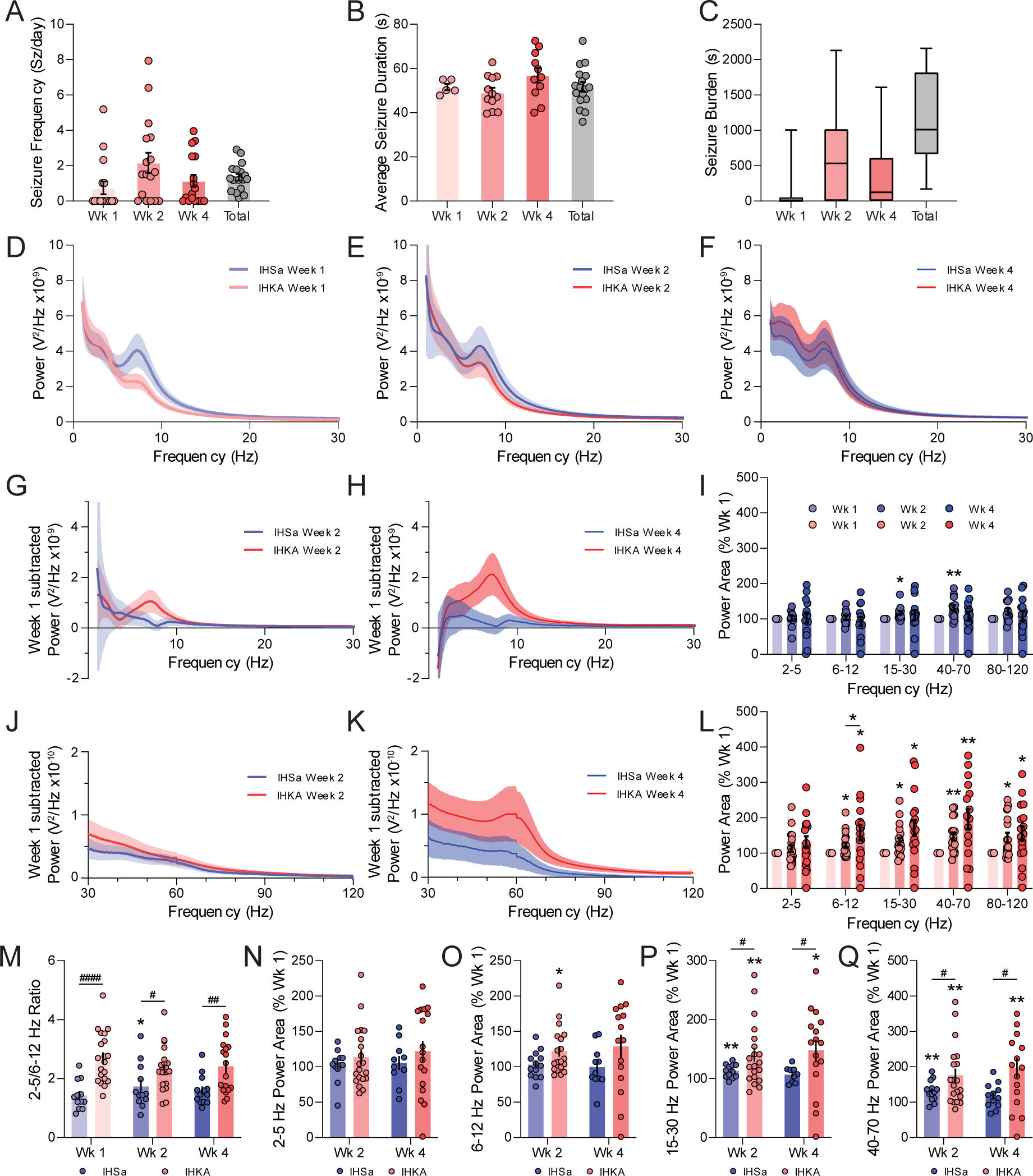
Intra-BLA LFP depth electrode recordings during development of SE. From a depth electrode placed within the BLA in mice injected with IHKA, measures including seizure frequency (A), average seizure duration (B), and seizure burden (C) were analyzed over a period of 4 weeks. During interictal periods in IHKA mice the LFP power spectra were analyzed across those same timepoints and compared to the power spectra of IHSa mice to examine the effects of epilepsy progression on basal power spectral properties in IHKA mice (D, E, F). To better visualize these changes, the power spectra were split into lower (G, H) and higher (J, K) frequency ranges from weeks 2 and 4 were normalized to their week 1 power. Those power area changes were binned into frequency bands in IHSa (I) and IHKA (L) mice normalized to their week 1 values, with IHKA mice exhibiting more significant changes across frequency bands. To compare changes to power area across weeks resulting from IHKA treatment compared to IHSA, direct comparisons were made within frequency bands with ratiometric comparisons (M) or with power normalized to their week one values (N, O, P, Q) with notable significant differences observed in 2-5/6-12 Hz ratio, 15-30 Hz, and 40-70 Hz bands. * denotes the degree of significance between weeks within a condition, # denotes the degree of significance between conditions. I, L, M-Q derive significance from mixed-effects analysis with Šídák’s multiple comparisons test, data underwent outlier removal using the ROUT method (Q=1%), G-L, N-Q values reported normalized to their Wk 1 values. Data shown as Mean±SEM except in C where data are displayed as box-and-whiskers plot with whiskers showing min and max to emphasize all IHKA mice had seizure burden.

### Deficits in PV interneurons is sufficient to alter BLA network and behavioral states

To assess whether the loss of PV-positive interneurons in the BLA contributes to the deficits in network and behavioral states, we injected PV-Cre mice with either AAV-mCherry (control) or AAV-Flex-DTA (Figure 5A) to ablate an equivalent number of PV interneurons to that observed in chronically epileptic mice and examined the impact on BLA network activity and behavioral states. There is a reduction in the number of PV-positive interneurons in the BLA of AAV-DTA mice (7.96±0.42 cells/section) compared to controls (18.90±1.07 cells/section) (Figure 5B; n=109-111 sections, 6 mice per experimental group; p<0.0001; unpaired 2-tail t-test), which can be appreciated in the representative images from control (Figure 5C) and AAV-DTA-injected mice (Figure 5D).

**Figure 5.**
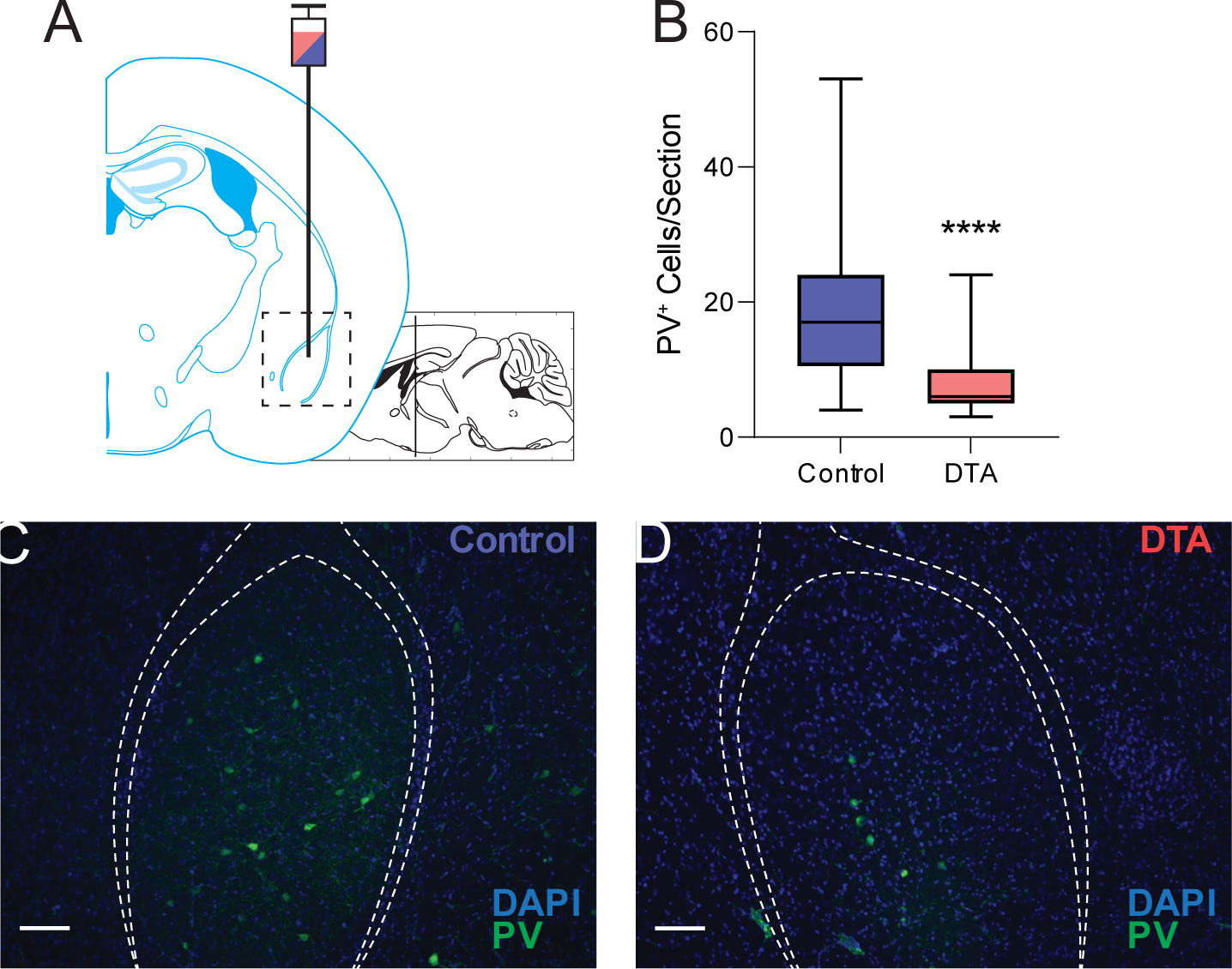
Genetically targeted viral ablation of PV^+^ interneurons in the BLA. A, Schematic showing the stereotaxic injection site of an AAV2-mCherry (control) or AAV-Flex-DTA virus into the BLA Adapted from Franklin & Paxinos (2008). B, DTA specifically ablates PV^+^ interneurons within the BLA to recapitulate the PV^+^ cell loss observed in the vIHKA model. C, D, Representative fluorescent IHC sections showing a decrease in the number of PV^+^ neurons in the BLA (white dashed outline) of DTA-injected mice compared to control virus injected mice. * denotes the degree of significance between conditions measured by unpaired 2-tail t-test. Data shown as a box-and-whiskers plot with whiskers showing min and max.

To examine whether the ablation of a large percentage of PV interneurons in the BLA impacts the function of remaining PV interneurons, we performed whole cell whole cell patch clamp recording and assessed the excitability and intrinsic membrane properties of the unablated PV interneurons. The input-output relationship in PV interneurons in the BLA is shifted in slices from AAV-DTA mice compared to controls (Figure 6A). There is a significant increase in the number of action potentials fired in response to a 100pA step in PV interneurons from AAV-DTA mice (19.4±2.5 Hz) compared to controls (7.5±4.3 Hz; p=0.020; unpaired 2-tail t-test), but not at 150 pA (Figure 6A; control: 22.0±7.2 Hz; DTA: 29.21±2.77; p=0.26; unpaired 2-tail t-test; n=7-16 cells, 3-5 mice per experimental group). The intrinsic membrane properties of PV interneurons in the BLA were compared between AAV-DTA mice and controls. There was a significant increase in the membrane time constant (control: 8.2±1.6 ms; DTA: 16.4±1.9 ms; p=0.012; unpaired 2-tail t-test) with no significant difference in membrane resistance (control: 401.4±71.0 MΩ; DTA: 432.8±44.0 MΩ; p=0.81; unpaired 2-tail t-test), rheobase (control: 81.3±12.2 pA; DTA: 56.4±7.2 pA; p=0.059; unpaired 2-tail t-test), or threshold (control: - 26.2±1.9 mV; DTA: -30.4±1.9 mV; p=0.17; unpaired 2-tail t-test) between control and AAV-DTA mice (Figure 6B; n=8-22 cells, 3-5 mice per experimental group).

**Figure 6.**
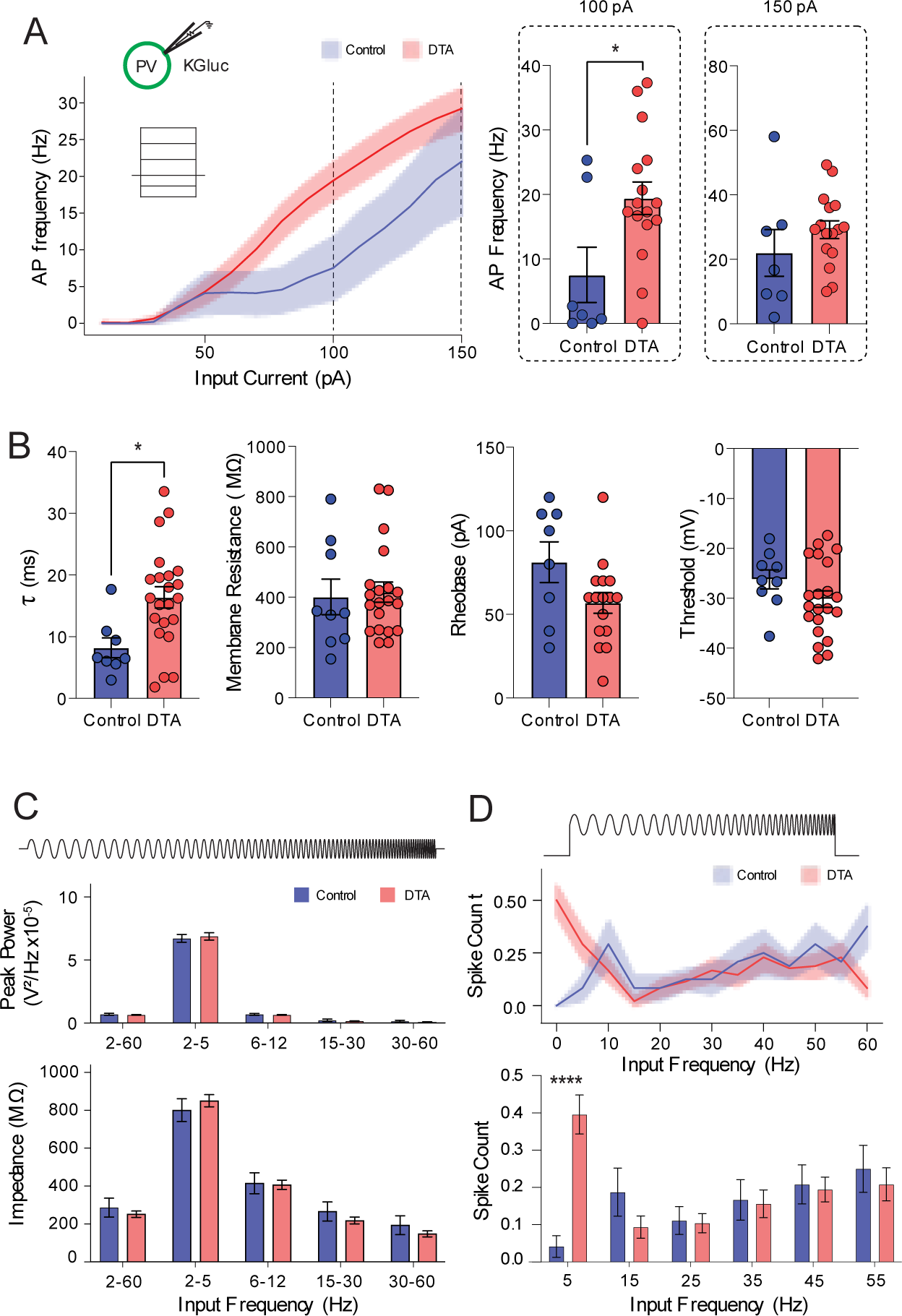
Partial loss of BLA PV+ interneurons shifts excitability and resonance. A, Schematic showing internal solution and genetically identified PV+ interneurons. DTA data represent remaining PV+ interneurons post-ablation. Input-output curves demonstrate that partial ablation of PV+ interneurons drives increased output in response to a 100 pA, but not a 150 pA, depolarizing square wave. B, The membrane time constant significantly increased in DTA mice (left), however other intrinsic membrane properties such as resistance (mid left), rheobase (mid right), and threshold (right) remain unaffected. C, There were no differences in peak power (top) or membrane impedance (bottom) between control and DTA mice. D, In response to a chirp stimulation consisting of an accelerating sine wave on top of a suprathreshold depolarizing current step, DTA mice demonstrated a significant treatment x frequency interaction with regard to spike generation. * denotes the degree of significance between conditions. The mean comparisons in A and B derive significance from unpaired 2-tailed t-tests. The two-factor comparisons in D derive significance from 2-way ANOVA with Šídák’s multiple comparisons test. Data shown as Mean±SEM.

The ablation of PV interneurons in the BLA did not alter the response of remaining PV interneurons to the subthreshold chirp stimulus. No differences were observed in the peak power in PV interneurons from AAV-DTA mice (1-60 Hz: 0.63±0.02 x10^-5^ V^2^/Hz; 2-5 Hz: 6.85±0.29 x10^-5^ V^2^/Hz; 6-12 Hz: 0.63±0.03 x10^-5^ V^2^/Hz; 15-30 Hz: 0.14±0.02 x10^-5^ V^2^/Hz; 30-60 Hz: 0.06±0.02 x10^-5^ V^2^/Hz) compared to controls (1-60 Hz: 0.68±0.08 x10^-5^ V^2^/Hz; 2-5 Hz: 6.70±0.32 x10^-5^ V^2^/Hz; 6-12 Hz: 0.67±0.08 x10^-5^ V^2^/Hz; 15-30 Hz: 0.21±0.09 x10^-5^ V^2^/Hz; 30-60 Hz: 0.13±0.08 x10^-5^ V^2^/Hz) (Figure 6C, top; Treatment effects: F_(1, 26)_ = 0.0107, p=0.92; 2-way ANOVA; n=8-20 cells, 3-5 mice per experimental group). Similarly, impedance was unchanged following DTA treatment (1-60 Hz: 253.1±16.6 MΩ; 2-5 Hz: 849.0±32.0 MΩ; 6-12 Hz: 407.2±24.7 MΩ; 15-30 Hz: 219.9±18.2 MΩ; 30-60 Hz: 149.8±16.0 MΩ) compared to controls (1-60 Hz: 287.4±49.7 MΩ; 2-5 Hz: 799.3±60.0 MΩ; 6-12 Hz: 415.3±55.3 MΩ; 15-30 Hz: 267.5±50.0 MΩ; 30-60 Hz: 195.0±49.9 MΩ) (Figure 6C; Treatment effects F_(1,26)_= 0.159, p=0.694; n=8-20 cells, 3-5 mice per experimental group). PV interneurons in the BLA of AAV-DTA mice exhibit an increase in the number of spikes elicited in response to the suprathreshold chirp stimulus at low input frequencies (DTA: 5 Hz: 0.40±0.05; 15 Hz: 0.09±0.03; 25 Hz: 0.10±0.03; 35 Hz: 0.16±0.04; 45 Hz: 0.19±0.03; 55 Hz: 0.21±0.04) compared to controls (5 Hz: 0.04±0.03; 15 Hz: 0.19±0.06; 25 Hz: 0.11±0.04; 35 Hz: 0.17±0.05; 45 Hz: 0.21±0.05; 55 Hz: 0.25±0.06) (Figure 6D; Interaction: F_(5, 132)_= 6.141, p<0.0001; 2-way ANOVA; n=8-16 cells, 3-5 mice per experimental group). These data suggest that the function of remaining PV interneurons and the ability to coordinate network states is impaired in mice with ablation of PV interneurons in the BLA.

The loss of PV interneurons in the BLA did not significantly alter the baseline BLA network activity (2-5 Hz: 2.57×10^-7^±6.17×10^-8^ V^2^; 6-12 Hz: 3.98×10-7±9.76×10-8 V^2^; 15-30 Hz: 1.19×10^-7^±2.68×10^-8^ V^2^; 40-70 Hz: 8.61×10^-8^±1.41×10^-8^ V^2^; 80-120 Hz: 2.22×10^-8^±3.60×10^-9^ V^2^; 2-5/6-12: 0.68±0.08) compared to controls (2-5 Hz: 1.59×10^-7^±4.07×10^-8^ V^2^; 6-12 Hz: 2.93×10^-7^±9.54×10^-8^ V^2^; 15-30 Hz: 9.47×10^-8^±2.94×10^-8^ V^2^; 40-70 Hz: 5.43×10^-8^±1.64×10^-8^ V^2^; 80-120 Hz: 1.31×10^-8^±3.17×10^-9^ V^2^; 2-5/6-12: 0.75±0.10) (Figure 7A, B; Frequency x Treatment Interaction: F_(4,65)_=0.38, p=0.820; 2-way ANOVA; Figure 7B inset: p=0.638; unpaired 2-tail t-test; N=7-8 mice per experimental group).

**Figure 7.**
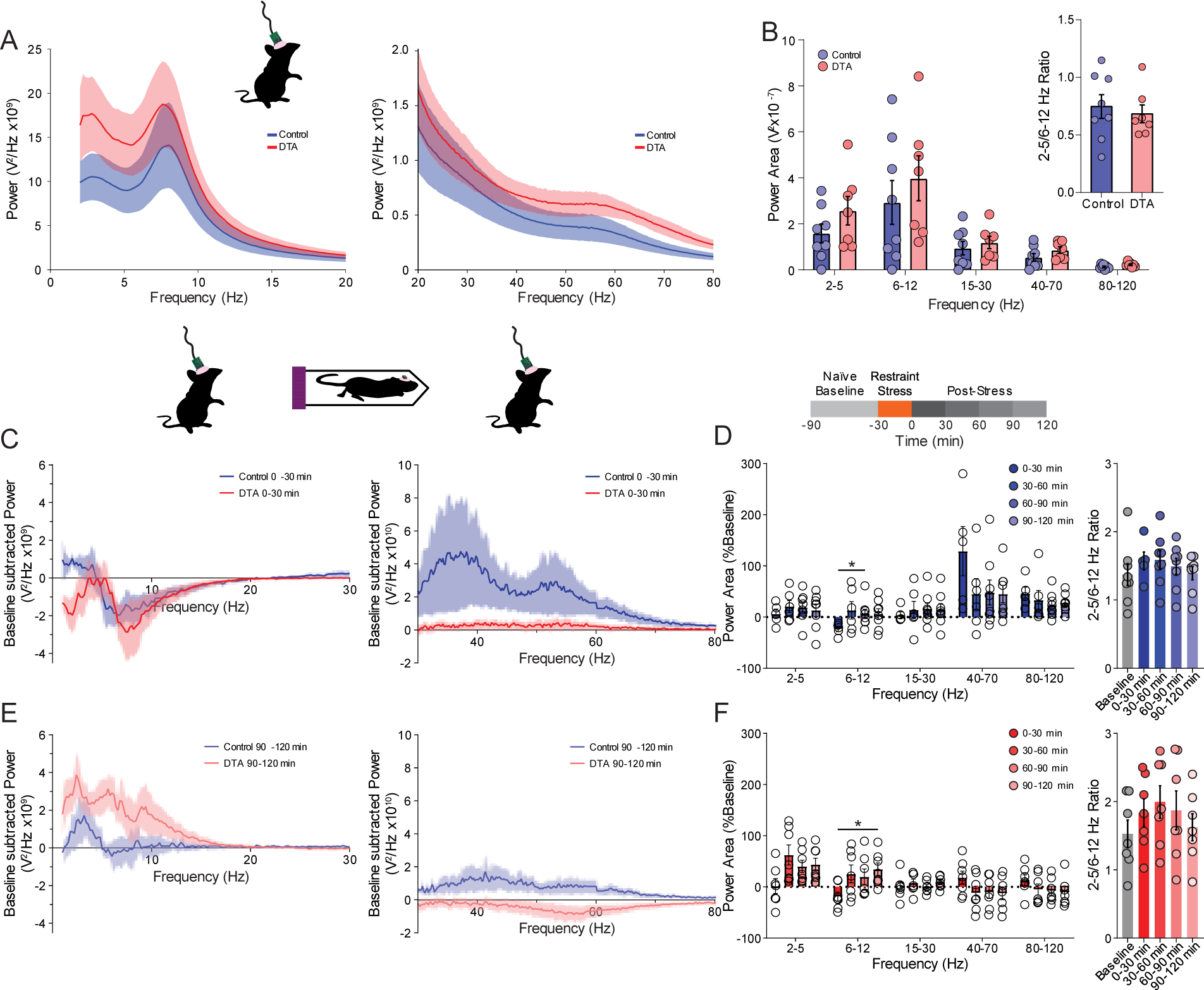
The loss of PV^+^ interneurons does not alter naïve or stressed network states in control and AAV-DTA mice. A, Power spectra from LFP recordings in the BLA of control and DTA injected mice showing an elevated basal power spectrum following the loss of PV^+^ interneurons in the BLA. B, However the increase in Power Area in unstressed mice was not significantly altered by the local, specific loss of PV^+^ interneurons. Further, the 2-5/6-12 Hz ratio (B, inset) remained unaltered between conditions at rest. To examine whether network oscillations during an active stress response were impaired, mice were exposed to an acute restraint stress, and the LFP was parsed into 30 minute segments to examine the temporal component of stress-induced changes at 0-30 minutes (C) and 90-120 minutes (E) following the acute stress. Control (D) and AAV-DTA (F) mice exhibited no significant stress-induced changes in the power area compared to their baseline values. * denotes significance within conditions between post-stress time bins, * denotes significance within conditions between post-stress time bins. B, inset derives significance from 2-tail t-test. B, F derives significance from 2-way ANOVA with Šídák’s multiple comparisons test, D derives significance from mixed-effects analysis with Šídák’s multiple comparisons test. Data shown as Mean±SEM.

With no network changes observed under basal conditions, an acute restraint stress was utilized to determine whether the loss of PV interneurons in the BLA would result in inappropriate network communication under stimulated conditions. A baseline, stress naïve LFP recording was taken prior to a 30 minute restraint stress to compare against the post-stress LFP recording at multiple post-stress timepoints. While some trends in the power area of pre-vs post-stress timepoints were observed within control (enhanced 40-70 and 80-120 Hz bands) and DTA (enhanced 2-5 Hz band) mice immediately (Figure 7C) and two hours following the stressor (Figure 7E) there were no significant changes in power compared to the baseline within control (Figure 7D: Frequency x Time Interaction: F_(16,115)_=2.891, p=0.0005; Mixed effects Analysis; 7D inset: Time effect: F_(1.464,9.152)_=2.859, p=0.117; Mixed-effect Analysis; N=5-8 mice) or AAV-DTA mice (Figure 7F: Frequency x Time Interaction: F_(16,120)_=5.793, p<0.0001; 2-way ANOVA; 7F inset: Time effect: F_(1.539,9.233)_=3.608, p=0.0775; 1-way ANOVA; N=7 mice). Repeated measures showed that in both control and DTA mice there were significant changes in the 6-12 Hz band 0-30 minutes following the acute restraint stress which then rebounded and became elevated for the remainder of the recording, reaching significance at 60-90 minutes in control and 90-120 minutes in DTA mice (Figure 7D, F). However, when the normalized power areas of control mice were directly compared with their time- and frequency-matched values in AAV-DTA mice, the loss of PV-interneurons in the BLA did not not significantly alter the LFP response to an acute restraint stress (direct comparison of Figure 7D, F not shown; Multiple unpaired t-tests with Holm-Šídák comparisons; N=5-8 mice per experimental group).

To examine changes in BLA network function following the specific loss of PV interneurons in a simplified, isolated preparation we carried out field recordings in the BLA of acute brain slices from control and AAV-DTA mice, examining the Power within a low and higher power band (Supplemental Figure 2). Here, we saw no significant change in the delta band power (Control: 0.026±0.011 μV^2^; DTA: 0.040±0.013 μV^2^; p=0.440; unpaired 2-tail t-test; n=13-17 slices, N= 4 mice per experimental group). However, we did observe a significant increase in AAV-DTA mice specific to the gamma band (Control: 0.034±0.006 μV^2^; DTA: 0.074±0.016 μV^2^; p=0.0473; unpaired 2-tail t-test; n=13-17 slices, N=4 mice per experimental group). Thus, these data consistently demonstrate in vivo and ex vivo the impact of PV interneuron loss in the BLA on coordination of gamma oscillations in the BLA.

These data suggest that the ability to coordinate oscillations in the BLA may be altered in mice with a loss of PV interneurons in the BLA. Given that these oscillatory states govern behavioral states, we examine the impact on avoidance behaviors and the behavioral expression of fear (Figure 8). AAV-DTA mice do not shown any difference in the amount of time spent in the center (89.4±9.3 s) or periphery (510.6±9.3 s) of the open field compared to controls (periphery: 514.0±7.5 s; center: 86.0±7.5 s) (Figure 8A; Treatment: F_(1,50)_=0.925, p=0.341; 2-way ANOVA; N=25-27 mice per experimental group). Similarly, AAV-DTA mice do not exhibit differences in the amount of time spent in the light (231.9±12.5 s) or dark (324.7±12.5 s) chamber of the light/dark box test compared to controls (light: 232.0±10.6 s; dark: 320.7±12.1 s) (Figure 8B; Treatment: F_(1,50)_=0.415, p=0.523; 2-way ANOVA; N=25-27 mice per experimental group). AAV-DTA mice do not exhibit differences in the amount of time spent in the closed (287.1±19.2 s), center (86.5±9.3 s), or open arms (121.6±14.3 s) of the elevated plus maze compared to controls (closed: 294.7±18.5 s; center: 71.7±6.5 s; open: 119.8±15.3 s) (Figure 8C; Treatment: F_(1,50)_=0.052, p=0.821; 2-way ANOVA; N=25-27 mice per experimental group). AAV-DTA also do not exhibit deficits in social interaction, spending an equivalent amount of time in the social (269.5±9.8 s), center (96.1±5.5 s), or empty chambers (215.2±9.6 s) compared to controls (social: 277.0±13.2 s; center: 92.7±4.5 s; empty: 206.0±12.1 s) (Figure 8D; Treatment: F_(1,50)_=2.49, p=0.121; 2-way ANOVA; N=25-27 mice per experimental group). These data suggest that the loss of PV interneurons in the BLA does not impact avoidance behaviors. However, the loss of PV interneurons in the BLA does alter the behavioral expression of fear. AAV-DTA mice exhibit a decrease in freezing during fear conditioning training (0.5: 2.8±1.0 %; 1.5: 2.4±0.8 %; 2.5: 7.6±1.9 %; 3.5: 18.2±2.7 %; 4.5: 28.4±3.5 %; 5.5: 30.5±3.9 %) compared to controls (0.5: 1.8±0.4 %; 1.5: 3.1±0.6 %; 2.5: 11.8±2.4 %; 3.5: 32.0±3.7 %; 4.5: 45.7±3.5 %; 5.5: 58.9±3.3 %) (Figure 8E; Treatment: F_(1,34)_=18.29, p=0.0001; Mixed-Effects Analysis; N=16-18 mice per experimental group). AAV-DTA mice also exhibit decreased freezing during contextual fear recall (3.5: 21.1±3.3 %; 4.5: 24.3±3.5 %) and cued fear recall (2.5: 22.6±2.8 %; 4.5: 16.5±1.9%) compared to controls (Contextual: 3.5: 46.0±4.5 %; 4.5: 50.4±5.2 %; Cued: 2.5: 37.7±4.5 %; 4.5: 42.0±5.7 %) (Figure 8F; Treatment: F_(1,34)_=15.08, p=0.0005; Mixed-Effects Analysis; Figure 7G: Treatment: F_(1,34)_=10.49, p=0.003; Mixed-Effects Analysis; N=17-18 mice per experimental group). These data demonstrate that the loss of PV interneurons in the BLA is sufficient to impair fear learning and recall of fear memories.

**Figure 8.**
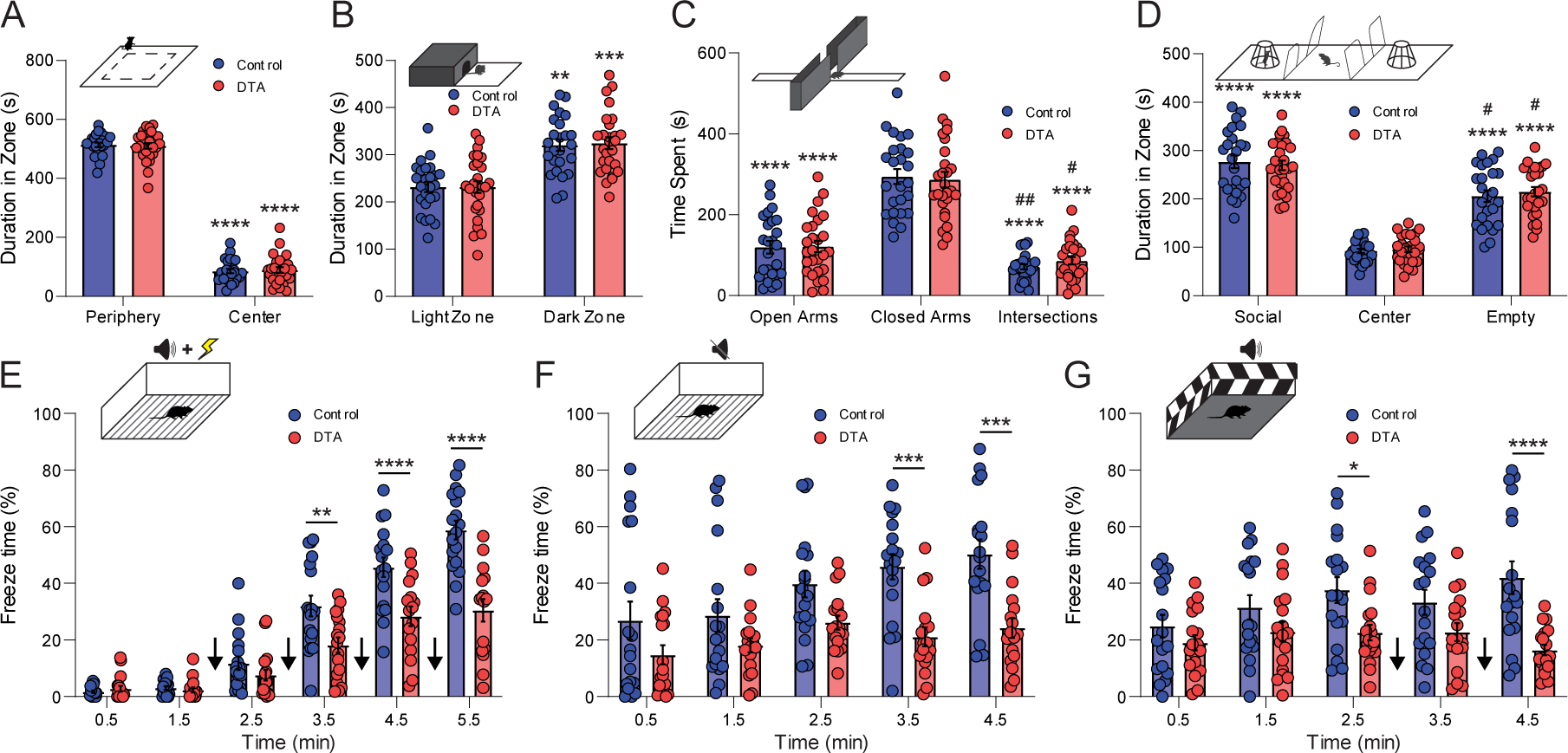
The specific loss of PV^+^ Interneurons impairs fear learning and recall of fear memories. A, Schematic of the open field test (top) in which both control and DTA mice spent significantly more time in the periphery of the open field, with no significant differences between treatment. B, Schematic of the light dark box (top) in which both control and DTA mice spent significantly less time in the light zone than the dark zone. C, Schematic of the elevated plus maze (top) in which control and DTA mice both spent significantly more time in the closed arms than either the open arms or intersection. D, Schematic of the 3-chamber social interaction test (top) in which no deficits in social interaction were observed between control and DTA mice. E, Schematic of the fear conditioning paradigm (top) in which control and mice were subjected to 4 bouts of tone-shock pairing (black arrows) while the time spent immobile was recorded. Control mice froze significantly more after the first tone-shock pairing. F, Schematic of the contextual recall paradigm (top) during which mice were placed into the fear conditioning chamber with the tone and shock disabled. Control mice froze longer than DTA mice across the time spent in the chamber, finally reaching significant levels in the final 2 minutes spent within the chamber. G, Schematic of the cued recall paradigm (top) in which the tone associated with the shock is played (black arrows) while in an environment different from the fear conditioning paradigm. Control mice exhibited significantly more freezing behavior prior to the first tone presentation and continued to freeze significantly more upon tone presentation. A, B: * denotes significance within condition compared to zone. C, D: * denotes significance compared to closed arms or center, # denotes significance compared to open arms or social zone. E, F, G: * denotes significance between conditions. A-D derives significance from 2-way ANOVA with Šídák’s multiple comparisons test. E-G derives significance from mixed-effects analysis with Šídák’s multiple comparisons test, data underwent outlier removal using the ROUT method (Q=1%). Data shown as Mean±SEM.

## Discussion

There is a need to better understand the mechanisms contributing to the well-established bidirectional pathophysiological relationship between psychiatric illnesses and epilepsy. This study examines whether the loss of PV interneurons in the BLA may contribute to behavioral deficits associated with epilepsy. Here we demonstrate profound behavioral deficits in chronically epileptic male C57Bl6/J mice (Figure 1) associated with a loss and dysfunction of PV interneurons in the BLA (Figures 2 and 3). Chronically epileptic mice also exhibit altered BLA network states (Figure 4) associated with the behavioral deficits. In an effort to directly determine whether deficits in PV interneurons in the BLA contributes to the network and behavioral deficits associated with chronic epilepsy, we used an AAV-Flex-DTA approach in PV-Cre mice to ablate an equivalent number of PV interneurons in the BLA to that observed in chronic epilepsy (Figure 5). We demonstrate that the loss of PV interneurons in the BLA is sufficient to induce behavioral deficits in fear learning and contextual- and cued-fear recall (Figure 8). These data demonstrate that the loss of PV interneurons in the BLA is capable of altering behavioral states, consistent with the critical role that PV interneurons in the BLA play orchestrating network states related to fear learning (Antonoudiou, 2020; Davis et al., 2017; Ozawa et al., 2020); however, the loss of PV interneurons in the BLA does not recapitulate the full spectrum of behavioral changes associated with chronic epilepsy, such as increased avoidance behaviors (compare Figures 1 and 8). Thus, it is not surprising that this pathophysiological mechanism may contribute to behavioral deficits associated with chronic epilepsy, but there are likely additional pathophysiological mechanisms at play.

The loss of PV interneurons in the BLA does not completely recapitulate the network and behavioral changes associated with chronic epilepsy which is not surprising given that there are other brain regions which are impacted in chronically epileptic animals, the most obvious of which is the hippocampus. There is well-documented interneuron loss in the hippocampus (Liu et al., 2014) which could also influence behavioral outcomes in this model. Further, it is likely that in the AAV-DTA model, the remaining PV interneurons in the BLA may be fully functional and able to compensate for the loss of PV interneurons; whereas, in the chronic epilepsy model, the remaining PV interneurons are dysfunctional (Figure 3). Further, other interneuron subtypes in the BLA are also impacted by epilepsy, but not AAV-DTA ablation. Therefore, ablating a subset of PV interneurons in the BLA does not fully recapitulate the neuropathological features of chronic epilepsy and, therefore, it is not surprising that this manipulation does not completely recapitulate the network and behavioral deficits observed in chronic epilepsy.

The current study focused on interneuron dysfunction given our existing knowledge regarding the role of interneurons in the BLA in driving behavioral states (Antonoudiou, 2020; Davis et al., 2017; Felix-Ortiz et al., 2016; Likhtik et al., 2014; Ozawa et al., 2020; Stujenske et al., 2014) (for review see (Tovote et al., 2015)). However, the data presented here also speaks to changes in excitatory synaptic inhibition. There is a shift in the cumulative distribution of the amplitude and frequency of sEPSCs on PV interneurons in the BLA in chronically epileptic mice (Figure 3). Further, there is an increase in the amplitude of sEPSCs and a shift in the cumulative distribution of the interevent interval of sEPSCs on principal neurons in the BLA (Supplemental Figure 1). Thus, it is clear that there are microcircuit changes beyond interneuron dysfunction that may contribute to the changes in network and behavioral states associated with chronic epilepsy.

Seizures are the most obvious manifestation of network dysfunction in epilepsy and, therefore, the primary focus of epilepsy research. We propose that more subtle deficits in network function may also underlie the risk for psychiatric illnesses comorbid with epilepsy. In fact, similar networks have been implicated in both epilepsy and psychiatric illnesses (Gilliam et al., 2004; Yang, 2018). Consistent with this theory, psychiatric illnesses are more common in patients with temporal lobe epilepsy (Kanner, 2017), suggesting that dysfunction in networks implicated in mood may increase vulnerability to psychiatric comorbidities. The majority of studies on temporal lobe epilepsy focus on the hippocampus. The current study suggests that other brain regions may also be altered in TLE models and may contribute to the larger symptom presentation associated with chronic epilepsy, such as psychiatric comorbidities.

The amygdala is known to play a critical role in emotional processing (LeDoux, 2000) and has been implicated in numerous psychiatric illnesses, with changes in activity correlating with symptom presentation (He et al., 2016; Li et al., 2015; Satterthwaite et al., 2016), outcome prediction (Connolly et al., 2017; Zhou et al., 2012), and treatment response (Cullen et al., 2016; Ellard et al., 2018; Fullana et al., 2017). Findings from the current study in mice suggest that the amygdala may also be involved in comorbid behavioral deficits and epilepsy. These data are supported by clinical studies demonstrating altered amygdala structure and enlargement associated with comorbid depression and epilepsy (Lv et al., 2014; Briellmann et al., 2007) and altered functional connectivity emanating from the amygdala in patients with temporal lobe epilepsy associated with psychiatric symptoms (Doucet et al., 2013). Amygdalar volume has been correlated with dysphoric disorders in epilepsy, including emotional instability, dysphoria, irritability, and aggression (Elst et al., 2009). Emerging and accumulating evidence points to amygdalar dysfunction in contributing to the full spectrum of symptoms in patients with temporal lobe epilepsy (Kullmann, 2011).

Given the evidence that the network communication within and between brain regions, such as the mPFC and BLA, drive specific behavioral states, particularly those with potential relevance to related to fear, anxiety, and depression (Antonoudiou, 2020; Davis et al., 2017; Felix-Ortiz et al., 2016; Likhtik et al., 2014; Ozawa et al., 2020; Stujenske et al., 2014) (for review see (Tovote et al., 2015)), it is necessary to explore how this network communication may become disrupted in epilepsy. Changes in functional connectivity involving the amygdala has been associated with psychiatric comorbidities in epilepsy (Colmers and Maguire, 2020; Yilmazer-Hanke et al., 2016). However, few studies have explored functional changes in network communication associated with comorbid psychiatric illnesses and epilepsy. Here we demonstrate that BLA network states are altered in chronically epileptic mice (Figure 4). We propose that the corruption in network activity within the BLA, and potentially between brain regions, contributes to the behavioral deficits associated with chronic epilepsy in mice. Further studies are required to fully understand changes in the network communication between brain regions and the impact on behavioral states associated with epilepsy. Further, the network communication driving behavioral states is dynamic (Antonoudiou, 2020; Davis et al., 2017; Felix-Ortiz et al., 2016; Likhtik et al., 2014; Ozawa et al., 2020; Stujenske et al., 2014) (for review see (Tovote et al., 2015)). The current study focused on changes in network states at baseline and did not explore dynamic changes associated with dynamic shifts in behavioral states. Thus, additional studies will be required to fully understand the relationship between network and behavioral states under pathological conditions, such as epilepsy.

This study investigates the cellular and circuit mechanisms contributing to comorbid behavioral deficits in epilepsy, pointing to a role for PV interneuron loss in the BLA and dysfunction in the network communication driving behavioral states. Despite the well-established role for network communication between brain regions driving behavioral states, this is the first study to our knowledge investigating changes in network communication under pathological conditions in contributing to comorbid behavioral deficits. These data demonstrate that altered network states in brain regions involved in emotional processing likely contribute to comorbid behavioral changes associated with chronic epilepsy.

**Supplemental Figure 1.**
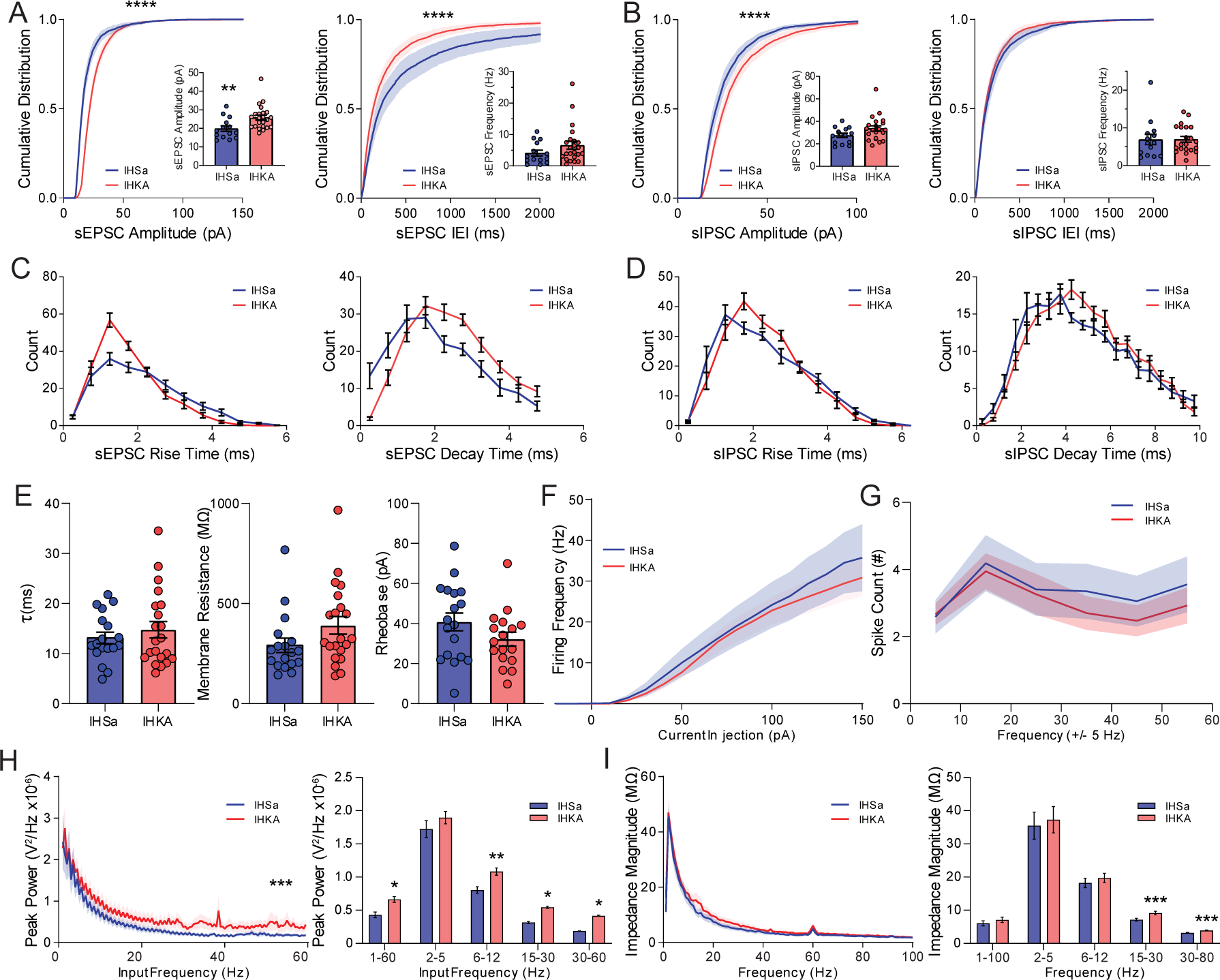
Active and passive electrophysiological properties of BLA Principal neurons. A. Whole-cell patch clamp recordings were taken from BLA principal neurons, and the cumulative distribution of sEPSCs in IHKA mice were observed to have a right-shift in amplitude and a left-shift in IEI, with a corresponding increase in mean amplitude, but no significant difference in mean frequency. B. BLA principal neurons showed a right-shift in the cumulative distribution of sIPSC amplitude in IHKA mice with no change in sIPSC IEI, with no significant changes in the mean amplitude or frequency of sIPSC afferents. Both sEPSCs and sIPSCs saw no significant changes in rise (C) or decay (D) kinetics between IHSa and IHKA mice. E, Intrinsic membrane properties including the membrane time constant (left), membrane resistance (middle), and rheobase (right) are not significantly altered in BLA principal neurons in IHKA mice. Firing properties including the input-output relationship (F) and firing in response to a supra-threshold chirp stimulation (G) did not reveal any significant differences between IHSa and IHKA mice. H, A subthreshold Chirp current (top) was injected into the current-clamped principal neuron to determine intrinsic passive resonant membrane properties at different input frequencies, IHKA principal neurons exhibited increased resonant properties across the wavelengths tested (left) and within specific high-frequency bands (right). I, Membrane impedance as a function of frequency was not significantly higher in IHKA across the entire frequency range (left) however there were some higher frequency bands that were significantly increased (right). * denotes the degree of significance between conditions. Cumulative distributions in A, B derive significance from Kolmogorov-Smirnov tests. The mean comparisons in A, B, and E derive significance from unpaired 2-tailed t-tests. Histograms in C, D derive significance from Wilcoxon matched pairs signed rank test, 2-tailed. The two-factor comparisons in F-I derive significance from 2-way ANOVA with Šídák’s multiple comparisons test. Data shown as Mean±SEM.

**Supplemental Figure 2.**
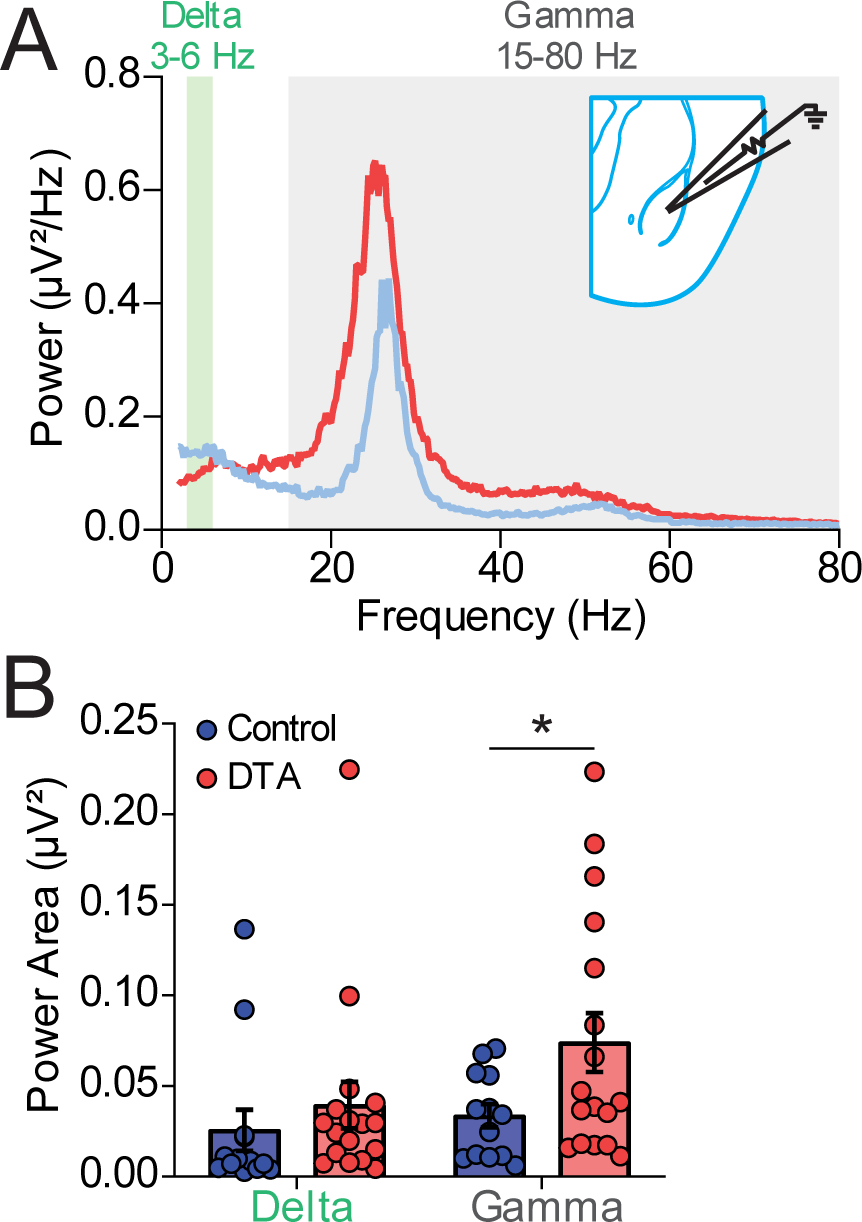
Specific loss of PV^+^ Interneurons alters local network synchronization. A. Representative local field potential power spectra recorded in the BLA of acute brain slices (inset schematic) taken from mice treated with DTA or control virus. Highlighted regions represent the frequency bands taken for the power area. B. Mean power area in Control and DTA injected slices showing no change in delta power with a significant increase in the gamma range following the specific loss of BLA PV interneurons. * denotes the degree of significance between conditions. The mean comparisons in B derive significance from unpaired 2-tailed t-tests. Data shown as Mean±SEM.

## Supporting information

Stats Table

## Author Contributions

PC and JM designed research; PC, TB, and GS performed research; PC and PA analyzed data; PC, PA, and JM wrote the paper. PF provided critical resources for this project.

