## Supplementary material for "Loss of PV interneurons in the BLA contributes to altered network and behavioral states in chronically epileptic mice": Stats Table

| Figure | Pannel | Experiment | Measure | | | | |
|---|---|---|---|---|---|---|---|
| Figure 1 | A | Open Field | 2-way ANOVA | vIHSa vs vIHKA | Interaction | **F(1,19)=17.10, p=0.0006** | |
| | | | | | Position | **F(1,19)=1132, p<0.0001** | |
| | | | | | Treatment | **F(1,19)=9.939, p=0.0052** | |
| | | | | vIHSa n=10 | vIHKA n=11 | | |
| | | | Periphery | 500.22±8.18s | 556.34±10.57s | | |
| | | | Center | 99.78±8.18s | 43.66±10.57s | | |
| | B | Light/Dark | 2-way ANOVA | vIHSa vs vIHKA | Interaction | **F(1,19)=57.99, p<0.0001** | |
| | | | | | Position | **F(1,19)=95.80, p<0.0001** | |
| | | | | | Treatment | **F(1,19)=13.11, p= 0.0018** | |
| | | | | vIHSa n=10 | vIHKA n=11 | | |
| | | | Light | 249.02±19.45s | 47.75±18.56s | | |
| | | | Dark | 310.39±20.35s | 539.38±21.86s | | |
| | C, left | EPM time in open | unpaired 2-tail t-test | vIHSa vs vIHKA | p=0.188 | t=1.367, df=19 | |
| | | | | vIHSa n=10 | vIHKA n=11 | | |
| | | | | 84.17±15.43s | 125.30±24.96s | | |
| | C, right | EPM open entries | unpaired 2-tail t-test | vIHSa vs vIHKA | **p=0.0193** | t=2.557, df=19 | |
| | | | | vIHSa n=10 | vIHKA n=11 | | |
| | | | | 45.60±9.27 entries | 21.00±3.67 entries | | |
| | D | Sucrose Preference | unpaired 2-tail t-test | vIHSa vs vIHKA | **p=0.0003** | t=4.357, df=19 | |
| | | | | vIHSa n=10 | vIHKA n=11 | | |
| | | | | 85.14±3.03% | 55.73±5.81% | | |
| | | | | PV+ Sa | PV+ KA | | |
| Figure 2 | B | Cell Count | vIHKA induced Cell loss | 21.52±0.95, n=122, N=8 | 13.60±0.73, n=120, N=8 | | |
| | | | unpaired 2-tail t-test | PV+Sa vs PV+KA | **p=<0.0001** | t=6.57, df=240 | |
| | | **sEPSC** | | PV+ Sa n=32, N=13 | PV+ KA, n=31, N=11 | PV- Sa, n=14, N=4 | PV- KA, n=23, N=5 |
| Figure 3 | | **Mean Amp** | | -31.04±2.39 pA, n=32 | -40.71±4.15 pA, n=31 | -19.85±1.50 pA, n=14 | -25.78±1.35 pA, n=23 |
| | A, inset left | | unpaired 2-tail t-test | PV+Sa vs PV+KA | **p=0.0462** | t=2.04, df=61 | |
| | -- | | unpaired 2-tail t-test | PV-Sa vs PV-KA | **p=0.0073** | t=2.85, df=35 | |
| | | **Amp Cumulative Distribution** | | | | | |
| | A, left | | Kolmogorov-Smirnov test | PV+Sa vs PV+KA | **p<0.0001** | | |
| | -- | | Kolmogorov-Smirnov test | PV-Sa vs PV-KA | **p<0.0001** | | |
| | | | | PV+ Sa n=32, N=13 | PV+ KA, n=31, N=11 | PV- Sa, n=14, N=4 | PV- KA, n=23, N=5 |
| | | **Mean Freq** | | 16.11±2.42 Hz, n=32 | 15.13±2.47 Hz, n=31 | 4.08±0.92 Hz, n=14 | 6.56±1.26 Hz, n=23 |
| | A, inset right | | unpaired 2-tail t-test | PV+Sa vs PV+KA | p=0.7759 | t=0.29, df=61 | |
| | -- | | unpaired 2-tail t-test | PV-Sa vs PV-KA | p=0.1701 | t=1.40, df=35 | |
| | | **Freq Cumulative Distribution** | | | | | |
| | A, right | | Kolmogorov-Smirnov test | PV+Sa vs PV+KA | **p<0.0001** | | |
| | -- | | Kolmogorov-Smirnov test | PV-Sa vs PV-KA | **p<0.0001** | | |
| | | **Rise** | | | | | |
| | C, left | | Wilcoxon matched pairs signed rank test, two-tailed | PV+Sa vs PV+KA | p=0.2334 | W=-32 | |
| | -- | | Wilcoxon matched pairs signed rank test, two-tailed | PV-Sa vs PV-KA | p=0.5186 | W=-18 | |
| | | **Decay** | | | | | |
| | C, right | | Wilcoxon matched pairs signed rank test, two-tailed | PV+Sa vs PV+KA | p>0.9999 | W=1 | |
| | -- | | Wilcoxon matched pairs signed rank test, two-tailed | PV-Sa vs PV-KA | p=0.5566 | W=13 | |
| | | **sIPSC** | | PV+ Sa n=32, N=13 | PV+ KA, n=31, N=11 | PV- Sa, n=14, N=4 | PV- KA, n=23, N=5 |
| | | **Mean Amp** | | 28.47±1.77 pA, n=32 | 28.63±1.23 pA, n=31 | 27.50±1.92 pA, n=14 | 33.62±2.43 pA, n=21 |
| | B, inset left | | unpaired 2-tail t-test | PV+Sa vs PV+KA | p=0.9391 | t=0.077, df=61 | |
| | -- | | unpaired 2-tail t-test | PV-Sa vs PV-KA | p=0.0780 | t=1.82, df=33 | |
| | | **Amp Cumulative Distribution** | | | | | |
| | B, left | | Kolmogorov-Smirnov test | PV+Sa vs PV+KA | p=0.1514 | | |
| | -- | | Kolmogorov-Smirnov test | PV-Sa vs PV-KA | **p<0.0001** | | |
| | | | | PV+ Sa n=32, N=13 | PV+ KA, n=31, N=11 | PV- Sa, n=14, N=4 | PV- KA, n=23, N=5 |
| | | **Mean Freq** | | 8.00±1.18 Hz, n=32 | 5.12±0.67 Hz, n=31 | 6.90±1.40 Hz, n=14 | 6.96±0.78 Hz, n=21 |
| | B, inset right | | unpaired 2-tail t-test | PV+Sa vs PV+KA | **p=0.0395** | t=2.10, df=61 | |
| | -- | | unpaired 2-tail t-test | PV-Sa vs PV-KA | p=0.9680 | t=0.040, df=33 | |
| | | **Freq Cumulative Distribution** | | | | | |
| | B, right | | Kolmogorov-Smirnov test | PV+Sa vs PV+KA | **p<0.0001** | | |
| | -- | | Kolmogorov-Smirnov test | PV-Sa vs PV-KA | p=0.1438 | | |
| | | **Rise** | | | | | |
| | D, left | | Wilcoxon matched pairs signed rank test, two-tailed | PV+Sa vs PV+KA | p>0.9999 | W=-1 | |
| | -- | | Wilcoxon matched pairs signed rank test, two-tailed | PV-Sa vs PV-KA | p=0.5693 | W=-16 | |
| | | **Decay** | | | | | |
| | D, right | | Wilcoxon matched pairs signed rank test, two-tailed | PV+Sa vs PV+KA | p=0.7285 | W=-20 | |
| | -- | | Wilcoxon matched pairs signed rank test, two-tailed | PV-Sa vs PV-KA | p=0.8194 | W=13 | |
| | | **Passive Properties** | | PV+ Sa n=17, N=5 | PV+ KA n=33 N=10 | PV- Sa, n=18, N=5 | PV- KA, n=21, N=7 |
| | | **Tau** | | 11.67±1.41 ms, n=17 | 9.11±0.69 ms, n=33 | 13.12±1.15 ms, n=18 | 14.77±1.66 ms, n=21 |
| | E, left | | unpaired 2-tail t-test | PV+Sa vs PV+KA | p=0.0716 | t=1.84, df=48 | |
| | -- | | unpaired 2-tail t-test | PV-Sa vs PV-KA | p=0.433 | t=0.79, df=37 | |
| | | | | PV+ Sa n=17, N=5 | PV+ KA n=33 N=10 | PV- Sa, n=18, N=5 | PV- KA, n=21, N=7 |
| | | **Rm** | | 432.6±51.4 MΩ, n=17 | 342.6±27.4 MΩ, n=33 | 291.3±35.7 MΩ, n=18 | 390.5±43.8 MΩ, n=21 |
| | E, middle | | unpaired 2-tail t-test | PV+Sa vs PV+KA | p=0.0955 | t=1.70, df=48 | |
| | -- | | unpaired 2-tail t-test | PV-Sa vs PV-KA | p=0.0942 | t=1.72, df=37 | |
| | | | | PV+ Sa, n=28, N=8 | PV+ KA, n=31, N=10 | PV- Sa, n=18, N=5 | PV- KA, n=17, N=7 |
| | | **Rheobase** | | 27.01±2.94 pA, n=28 | 30.91±3.21 pA, n=31 | 40.84±4.49 pA, n=18 | 32.20±3.44 pA, n=17 |
| | E, right | | unpaired 2-tail t-test | PV+Sa vs PV+KA | p=0.377 | t=0.891, df=57 | |
| | -- | | unpaired 2-tail t-test | PV-Sa vs PV-KA | p=0.139 | t=1.52, df = 33 | |
| | | **IO** | | | | | |
| | F | | 2way ANOVA | PV+Sa vs PV+KA | Interaction | F(16,913)=0.80, p=0.692 | |
| | | | | | Current Step | **F(16,913)=44.76, p<0.0001** | |
| | | | | | **Treatment** | **F(1,913)=9.39, p=0.0022** | |
| | -- | | 2way ANOVA | PV-Sa vs PV-KA | Interaction | F(16,612)=0.087, p>0.9999 | |
| | | | | | Current Step | **F(16,612)=20.38, p<0.0001** | |

| | | | | | | | |
|---|---|---|---|---|---|---|---|
| | | | | | **Treatment** | F(1,612)=1.90, p=0.169 | |
| | | **Short Chirp** | | | | | |
| | **G** | | 2way ANOVA | PV+Sa vs PV+KA | Interaction | F(5,276)=0.149, p=0.980 | |
| | | | | | Frequency | F(5,276)=1.29, p=0.267 | |
| | | | | | **Treatment** | F(1,276)=2.88, p=0.0906 | |
| | **--** | | 2way ANOVA | PV-Sa vs PV-KA | Interaction | F(5,210)=0.111, p=0.990 | |
| | | | | | Frequency | F(5,210)=1.28, p=0.272 | |
| | | | | | **Treatment** | F(1,210)=0.956, p=0.329 | |
| | | **Chirp Peak Power** | | | | | |
| | **H, left** | | 2way ANOVA | PV+Sa vs PV+KA | Interaction | **F(4,290)=42.16 p<0.0001** | |
| | | | | | Frequency | **F(4,290)=217.5, p<0.0001** | |
| | | | | | **Treatment** | **F(1,290)=232.3, p<0.0001** | |
| | **--** | | 2way ANOVA | PV-Sa vs PV-KA | Interaction | F(4,180)=0.209, p=0.933 | |
| | | | | | Frequency | **F(4,180)=219.8, p<0.0001** | |
| | | | | | **Treatment** | **F(1,180)=38.57, p<0.0001** | |
| | | | | PV+ Sa, n=27, N=7 | PV+ KA, n=33, N=10 | PV- Sa, n=18, N=5 | PV- KA, n=20, N=7 |
| | **H, right** | | 1-60 Hz | $1.72x10^{-6}\pm 1.55x10^{-7}$ | $8.05x10^{-7}\pm 6.30x10^{-8}$ | $4.31x10^{-7}\pm4.24x10^{-8}$ | $6.63x10^{-7}\pm4.24x10^{-8}$ |
| | | | 2-5 Hz | $6.57x10^{-6}\pm3.99x10^{-7}$ | $2.68x10^{-6}\pm 1.66x10^{-7}$ | $1.72x10^{-6}\pm1.27x10^{-7}$ | $1.90x10^{-6}\pm9.35x10^{-8}$ |
| | | | 6-12 Hz | $3.00x10^{-6}\pm2.21x10^{-7}$ | $1.34x10^{-6}\pm 7.89x10^{-8}$ | $8.04x10^{-7}\pm4.90x10^{-8}$ | $1.08x10^{-6}\pm5.39x10^{-8}$ |
| | | | 15-30 Hz | $1.08x10^{-6}\pm4.22x10^{-8}$ | $5.90x10^{-7}\pm 3.04x10^{-8}$ | $3.16x10^{-7}\pm1.38x10^{-8}$ | $5.45x10^{-7}\pm1.49x10^{-8}$ |
| | | | 30-60 Hz | $9.78x10^{-7}\pm4.07x10^{-8}$ | $4.74x10^{-7}\pm 1.68x10^{-8}$ | $1.88x10^{-7}\pm2.92x10^{-9}$ | $4.17x10^{-7}\pm9.05x10^{-9}$ |
| | | | Multiple unpaired 2-tail t-test | PV+Sa vs PV+KA | 1-60 Hz | **p<0.001** | t=5.87, df=58 |
| | | | | | 2-5 Hz | **p<0.001** | t=9.60, df=58 |
| | | | | | 6-12 Hz | **p<0.001** | t=7.67, df=58 |
| | | | | | 15-30 Hz | **p<0.001** | t=9.64, df=58 |
| | | | | | 30-60 Hz | **p<0.001** | t=12.24, df=58 |
| | **--** | | Multiple unpaired 2-tail t-test | PV-Sa vs PV-KA | 1-60 Hz | **p=0.001** | t=3.85, df=36 |
| | | | | | 2-5 Hz | p=0.274 | t=1.11, df=36 |
| | | | | | 6-12 Hz | **p=0.001** | t=3.80, df=36 |
| | | | | | 15-30 Hz | **p<0.001** | t=11.25, df=36 |
| | | | | | 30-60 Hz | **p<0.001** | t=22.96, df=36 |
| | | **Reverse Chirp Peak Power** | | | | adjusted p value | |
| Removed | **Not shown** | | 2way ANOVA | PV+Sa Ch vs PV+Sa rCh | Interaction | F(4,100)=0.45, p=0.772 | |
| | | | | | Frequency | **F(4,100)=213.2, p<0.0001** | |
| | | | | | **Chirp Direction** | F(1,100)=0.116, p=0.734 | |
| Removed | **Not shown** | | 2way ANOVA | PV+KA Ch vs PV+KA rCh | Interaction | **F(4,40)=7.27, p=0.0002** | |
| | | | | | Frequency | **F(4,40)=69.87, p<0.0001** | |
| | | | | | **Chirp Direction** | **F(1,40)=23.63, p<0.0001** | |
| | **Not shown** | | Holm-Šídák's multiple comparisons test | PV+Sa Ch vs PV+Sa rCh | 1-60 Hz | p=0.985 | t=0.083, df=20 |
| | | | alpha = 0.05 | | 2-5 Hz | p=0.928 | t=0.56, df=20 |
| | | | | | 6-12 Hz | p=0.985 | t=0.16, df=20 |
| | | | | | 15-30 Hz | p=0.158 | t=2.28, df=20 |
| | | | | | 30-60 Hz | p=0.123 | t=2.41, df=20 |
| | **Not shown** | | Holm-Šídák's multiple comparisons test | PV+KA Ch vs PV+KA rCh | 1-60 Hz | p=0.092 | t=2.34, df=8 |
| | | | alpha = 0.05 | | 2-5 Hz | **p=0.038** | t=3.19, df=8 |
| | | | | | 6-12 Hz | **p=0.004** | t=4.98, df=8 |
| | | | | | 15-30 Hz | p=0.092 | t=1.96, df=8 |
| | | | | | 30-60 Hz | **p=0.001** | t=6.18, df=8 |
| | | **Chirp Impedance** | | | | | |
| | **I, left** | | 2way ANOVA | PV+Sa vs PV+KA | Interaction | F(4,290)=0.18, p=0.947 | |
| | | | | | Frequency | **F(4,290)=111.6, p<0.0001** | |
| | | | | | **Treatment** | **F(1,290)=13.52, p=0.0003** | |
| | **S1, left** | | 2way ANOVA | PV-Sa vs PV-KA | Interaction | F(4,180)=0.036, p=0.998 | |
| | | | | | Frequency | **F(4,180)=94.36, p<0.0001** | |
| | | | | | **Treatment** | F(1,180)=1.29, p=0.258 | |
| | | | | PV+ Sa, n=27, N=7 | PV+ KA, n=33, N=10 | PV- Sa, n=18, N=5 | PV- KA, n=20, N=7 |
| | **I, right** | | 1-100 Hz | 9.60±1.00 MΩ | 13.92±1.10 MΩ | 6.08±0.75 | 7.10±0.80 |
| | | | 2-5 Hz | 47.20±4.72 MΩ | 53.75±4.62 MΩ | 35.48±4.08 | 37.28±3.96 |
| | | | 6-12 Hz | 26.40±1.88 MΩ | 33.56±1.98 MΩ | 18.24±1.42 | 19.74±1.42 |
| | | | 15-30 Hz | 12.19±0.66 MΩ | 17.72±0.75 MΩ | 7.16±0.45 | 9.13±0.54 |
| | | | 30-80 Hz | 5.55±0.18 MΩ | 9.36±0.21 MΩ | 3.20±0.11 | 3.92±0.15 |
| | | | Holm-Šídák's multiple comparisons test | PV+Sa vs PV+KA | 1-100 Hz | **p=0.018** | t=2.85, df=58 |
| | | | alpha = 0.05 | | 2-5 Hz | p=0.329 | t=0.98, df=58 |
| | | | | | 6-12 Hz | **p=0.025** | t=2.58, df=58 |
| | | | | | 15-30 Hz | **p<0.001** | t=5.39, df=58 |
| | | | | | 30-80 Hz | **p<0.001** | t=13.43, df=58 |
| | **--** | | Holm-Šídák's multiple comparisons test | PV-Sa vs PV-KA | 1-100 Hz | p=0.74 | t=0.93, df=36 |
| | | | alpha = 0.05 | | 2-5 Hz | p=0.75 | t=0.32, df=36 |
| | | | | | 6-12 Hz | p=0.74 | t=0.75, df=36 |
| | | | | | 15-30 Hz | **p=0.036** | t=2.76, df=36 |
| | | | | | 30-80 Hz | **p=0.003** | t=3.81, df=36 |
| | | **Reverse Chirp Impedance** | | | | | |
| | **Not shown** | | 2way ANOVA | PV+Sa Ch vs PV+Sa rCh | Interaction | F(4,100)=0.14, p=0.968 | |
| | | | | | Frequency | **F(4,100)=142.3, p<0.0001** | |
| | | | | | **Treatment** | F(1,100)=0.98, p=0.324 | |
| | **Not shown** | | 2way ANOVA | PV+KA Ch vs PV+KA rCh | Interaction | F(4,40)=0.008, p>0.9999 | |
| | | | | | Frequency | **F(4,40)=49.80, p<0.0001** | |
| | | | | | **Treatment** | F(1,40)=0.073, p=0.789 | |
| | **Not shown** | | Holm-Šídák's multiple comparisons test | PV+Sa Ch vs PV+Sa rCh | 1-100 Hz | p=0.905 | t=0.62, df=20 |
| | | | alpha = 0.05 | | 2-5 Hz | p=0.905 | t=0.27, df=20 |
| | | | | | 6-12 Hz | p=0.858 | t=0.89, df=20 |
| | | | | | 15-30 Hz | p=0.905 | t=0.51, df=20 |
| | | | | | 30-80 Hz | p=0.852 | t=1.02, df=20 |
| | **Not shown** | | Holm-Šídák's multiple comparisons test | PV+KA Ch vs PV+KA rCh | 1-100 Hz | p=0.994 | t=0.05, df=8 |
| | | | alpha = 0.05 | | 2-5 Hz | p=0.994 | t=0.12, df=8 |
| | | | | | 6-12 Hz | p=0.994 | t=0.24, df=8 |
| | | | | | 15-30 Hz | p=0.984 | t=0.48, df=8 |
| | | | | | 30-80 Hz | p=0.051 | t=3.33, df=8 |
| **Figure 4** | **A** | **Sz Frequency** | 1way ANOVA | KA Sz Wk 1, 2, 4, total | **Treatment** | F(3,60)=2.48, p=0.070 | |
| | | | N=16 | Wk1 | Wk2 | Wk4 | Total |
| | | | | 0.79±0.38 | 2.30±0.59 | 1.22±0.36 | 1.39±0.20 |
| | **B** | **Sz Duration** | 1way ANOVA | KA Sz Wk 1, 2, 4, total | **Treatment** | F(3,40)=1.66, p=0.19 | |

N=5-12

| Wk1 | Wk2 | Wk4 | Total |
|---|---|---|---|
| 51.61±1.45 s | 49.11±2.17 s | 56.91±3.23 s | 52.60±2.08 s |

**C** **Sz Burden** 1way ANOVA KA Sz Wk 1, 2, 4, total **Treatment** **F(2.036,30.54)=14.45, p<0.0001**

N=16

| Wk1 | Wk2 | Wk4 | Total |
|---|---|---|---|
| 105.94±65.95 s | 653.12±151.04 s | 381.56±114.90 s | 1140.63±161.11 s |

**I** **Power Area Sa** Mixed-effects analysis BLA SA Sz Wk 1, 2, 4

| | | |
|---|---|---|
| Frequency | F(4,65)=1.23, p=0.308 | (outliers removed) |
| Week | **F(1.34,81.99)=3.85, p=0.041** | N=12-14 |
| **Interaction** | F(8,122)=0.62, p=0.764 | |

| | BLA Sa Wk 2 | BLA Sa Wk 4 | Adjusted p values Wk1 v Wk2 | Wk1 v Wk4 | Wk2 v Wk4 |
|---|---|---|---|---|---|
| 2-5 | 101.8±6.6 | 102.8±15.2 | 0.9545 | 0.9811 | 0.9964 |
| 6-12 | 103.1±5.3 | 93.2±12.4 | 0.8231 | 0.8482 | 0.6671 |
| 15-30 | 117.3±5.3 | 111.0±13.7 | **0.0149** | 0.7058 | 0.8932 |
| 40-70 | 131.3±8.0 | 108.5±12.8 | **0.0049** | 0.7833 | 0.3742 |
| 80-120 | 121.1±8.8 | 108.5±14.2 | 0.073 | 0.8238 | 0.7685 |

**L** **Power Area KA** Mixed-effects analysis BLA KA Week 1,2,4

| | | |
|---|---|---|
| Frequency | F(4,95)=1.65, p=0.168 | (outliers removed) |
| Week | **F(1.35,114.8)=37.14, p<0.0001** | N=16-20 |
| **Interaction** | F(8,170)=1.21, p=0.298 | |

| | BLA KA Wk 2 | BLA KA Wk 4 | Adjusted p values Wk1 v Wk2 | Wk1 v Wk4 | Wk2 v Wk4 |
|---|---|---|---|---|---|
| 2-5 | 113.7±9.7 | 131.2±16.6 | 0.353 | 0.1605 | 0.4225 |
| 6-12 | 122.1±7.7 | 158.5±21.9 | **0.0236** | **0.0355** | **0.0401** |
| 15-30 | 133.0±10.1 | 170.8±22.3 | **0.0106** | **0.0123** | 0.0757 |
| 40-70 | 146.7±11.6 | 196.9±27.7 | **0.0019** | **0.0058** | 0.0563 |
| 80-120 | 143.7±13.7 | 156.0±20.6 | **0.0116** | **0.0309** | 0.6898 |

-- Power Sa v KA 3way ANOVA BLA SA KA Week 2,4

**M** **2-5/6-12 Ratio** Mixed-effects analysis BLA Sa vs KA week 1, 2, 4

| | |
|---|---|
| Week | F(1.68,48.77)=0.35, p=0.668 |
| Treatment | **F(1,31)=13.26, p=0.001** |
| **Interaction** | **F(2,58)=3.80, p=0.028** |

(outliers removed) N=12-20

| | BLA Sa | BLA KA |
|---|---|---|
| Wk 1 | 1.43±0.14 | 2.70±0.19 |
| Wk 2 | 1.73±0.22 | 2.35±0.17 |
| Wk 4 | 1.60±0.14 | 2.42±0.21 |

**N** **2-5 Power Area** Mixed-effects analysis BLA Sa vs KA week 1, 2, 4

| | |
|---|---|
| Week | F(1.65,45.26)=1.26, p=0.288 |
| Treatment | F(1,32)=1.45, p=0.237 |
| **Interaction** | F(2,55)=0.52, p=0.595 |

(outliers removed) N=10-20

| | BLA Sa | BLA KA |
|---|---|---|
| Wk 2 | 101.85±6.59 | 113.71±9.69 |
| Wk 4 | 105.46±9.90 | 122.10±14.70 |

**O** **6-12 Power Area** Mixed-effects analysis BLA Sa vs KA week 1, 2, 4

| | |
|---|---|
| Week | F(1.49,40.23)=2.26, p=0.130 |
| Treatment | **F(1,32)=4.63, p=0.039** |
| **Interaction** | F(2,54)=2.04, p=0.140 |

(outliers removed) N=11-20

| | BLA Sa | BLA KA |
|---|---|---|
| Wk 2 | 103.11±5.28 | 122.08±7.66 |
| Wk 4 | 99.56±8.77 | 129.04±16.33 |

**P** **15-30 Power Area** Mixed-effects analysis BLA Sa vs KA week 1, 2, 4

| | |
|---|---|
| Week | **F(1.77,46.91)=5.67; p=0.0082** |
| Treatment | **F(1,32)=5.89; p=0.021** |
| **Interaction** | F(2,53)=2.56; p=0.087 |

(outliers removed) N=9-20

| | BLA Sa | BLA KA |
|---|---|---|
| Wk 2 | 112.97±3.28 | 140.17±11.98 |
| Wk 4 | 106.69±4.45 | 147.90±17.89 |

**Q** **40-70 Power Area** Mixed-effects analysis BLA Sa vs KA week 1, 2, 4

| | |
|---|---|
| Week | **F(1.88,53.51)=11.28, p=0.0001** |
| Treatment | **F(1,32)=7.39, p=0.011** |
| **Interaction** | **F(2,57)=4.34, p=0.018** |

(outliers removed) N=12-20

| | BLA Sa | BLA KA |
|---|---|---|
| Wk 2 | 131.33±8.04 | 175.90±19.02 |
| Wk 4 | 117.42±10.02 | 196.93±27.68 |

**not shown** **80-120 Power Area** Mixed-effects analysis BLA Sa vs KA week 1, 2, 4

| | |
|---|---|
| Week | **F(1.79,51.78)=6.37; p=0.0045** |
| Treatment | F(1,32)=3.94; p=0.0558 |
| **Interaction** | F(2,58)=2.33; p=0.106 |

(outliers removed) N=12-20

| | BLA Sa | BLA KA |
|---|---|---|
| Wk 2 | 121.11±8.84 | 150.48±14.58 |
| Wk 4 | 108.47±14.17 | 155.96±20.59 |

**Figure 5** **B** **Cell Count** DTA induced Cell Loss

| Control | DTA | |
|---|---|---|
| 18.90±1.07, n=109, N=6 | 7.96±0.42, n=111, N=6 | |
| Control vs DTA | **p<0.0001** | t=9.57, df=218 |

unpaired 2-tail t-test

**Figure 6** **A** **I/O**

| | Control n=7 N=3 | DTA n=16 N=5 | unpaired 2-tail t-test | |
|---|---|---|---|---|
| 100 pA | 7.52±4.28 | 19.42±2.52 | **0.0204** | t=2.509, df=21 |
| 150 pA | 22±7.21 | 29.21±2.77 | 0.2637 | t=1.148, df=21 |

**B** **tau**

| Control n=8 N=3 | DTA n=22 N=5 | unpaired 2-tail t-test | |
|---|---|---|---|
| 8.22±1.6 ms | 16.39±1.94 ms | **0.0123** | t=2.676, df=28 |

**Rm**

| Control n=9 N=3 | DTA n=20 N=5 | unpaired 2-tail t-test | |
|---|---|---|---|
| 401.37±71.01 MΩ | 432.82±44.04 MΩ | 0.8058 | t=0.2483, df=27 |

**Rheobase**

| Control n=8 N=3 | DTA n=16 N=5 | unpaired 2-tail t-test | |
|---|---|---|---|
| 81.25±12.17 pA | 56.43±7.16 pA | 0.0593 | t=1.988, df=22 |

| | | | | | | |
|---|---|---|---|---|---|---|
| | **Threshold** | | Control n=9 N=3 | DTA n=21 N=5 | unpaired 2-tail t-test | |
| | | | -26.21±1.9 mV | -30.43±1.93 mV | 0.1702 | t=1.408, df=28 |

**Chirp Peak Power**

| | | | | | |
|---|---|---|---|---|---|
| **C, top** | 2way ANOVA | Control vs DTA | Interaction | F (4, 104) = 0.1887 | P=0.9438 |
| | | | Frequency | F (1.003, 26.08) = 669.1 | **P<0.0001** |
| | | | **Treatment** | F (1, 26) = 0.01070 | P=0.9184 |

| | Control n=8 N=3 | DTA n=20 N=5 | Holm-Šídák's multiple comparisons test alpha = 0.05 | |
|---|---|---|---|---|
| 1-60 Hz | 0.68±0.08x10-5 V2/Hz | 0.63±0.02x10-5 V2/Hz | 0.9877 | tf=0.57, df=8.36 |
| 2-5 Hz | 6.7±0.32x10-5 V2/Hz | 6.85±0.29x10-5 V2/Hz | 0.9982 | tf=0.37, df=18.98 |
| 6-12 Hz | 0.67±0.08x10-5 V2/Hz | 0.63±0.03x10-5 V2/Hz | 0.9952 | tf=0.46, df=8.66 |
| 15-30 Hz | 0.21±0.09x10-5 V2/Hz | 0.14±0.02x10-5 V2/Hz | 0.9558 | tf=0.77, df=7.74 |
| 30-60 Hz | 0.13±0.08x10-5 V2/Hz | 0.06±0.02x10-5 V2/Hz | 0.934 | tf=0.85, df=7.73 |

**Chirp Impedance**

| | | | | | |
|---|---|---|---|---|---|
| **C, bottom** | 2way ANOVA | Control vs DTA | Interaction | F (4, 104) = 2.644 | **P=0.0377** |
| | | | Frequency | F (1.096, 28.49) = 438.3 | **P<0.0001** |
| | | | **Treatment** | F (1, 26) = 0.1585 | P=0.6938 |

| | Control n=8 N=3 | DTA n=20 N=5 | Holm-Šídák's multiple comparisons test alpha = 0.05 | |
|---|---|---|---|---|
| 1-100 Hz | 287.45±49.7 | 253.1±16.63 | 0.9769 | tf=0.66, df=8.62 |
| 2-5 Hz | 799.27±60 | 849.01±32.05 | 0.9618 | tf=0.73, df=11.23 |
| 6-12 Hz | 415.35±55.32 | 407.16±24.66 | >0.9999 | tf=0.14, df=9.91 |
| 15-30 Hz | 267.46±50.01 | 219.94±18.24 | 0.9193 | tf=0.89, df=8.93 |
| 30-80 Hz | 195.02±49.87 | 149.78±15.98 | 0.9294 | tf=0.86, df=8.48 |

**Short Chirp**

| | | | | | |
|---|---|---|---|---|---|
| **D, top** | 2way ANOVA | Control vs DTA | Interaction | F (5, 132) = 6.141 | **P<0.0001** |
| | | | Frequency | F (5, 132) = 2.153 | P=0.0631 |
| | | | **Treatment** | F (1, 132) = 1.379 | P=0.2423 |

| | | Control n=8 N=3 | DTA n=16 N=5 | Holm-Šídák's multiple comparisons test alpha = 0.05 | |
|---|---|---|---|---|---|
| **D, bottom** | 0-10 Hz | 0.04±0.03 | 0.4±0.05 | **<0.0001** | tf=5.43, df=132 |
| | 10-20 Hz | 0.19±0.06 | 0.09±0.03 | 0.63 | tf=1.44, df=132 |
| | 20-30 Hz | 0.11±0.04 | 0.1±0.03 | >0.9999 | tf=0.11, df=132 |
| | 30-40 Hz | 0.17±0.05 | 0.16±0.04 | >0.9999 | tf=0.16, df=132 |
| | 40-50 Hz | 0.21±0.05 | 0.19±0.03 | >0.9999 | tf=0.21, df=132 |
| | 50-60 Hz | 0.25±0.06 | 0.21±0.04 | 0.9883 | tf=0.64, df=132 |

| | | | | | | |
|---|---|---|---|---|---|---|
| **Figure 7** | **B** | **Power Area**<br>N=7-8 | 2way ANOVA | Control vs DTA Basal EEG | **Interaction** | F(4,65)=0.383, p=0.820 |
| | | | | | Frequency | **F(4,65)=12.93, p<0.0001** |
| | | | | | Condition | F(1,65)=2.765, p=0.101 |

| | Ctrl | DTA |
|---|---|---|
| 2-5 | $1.59x10^{-7}$±$4.07x10^{-8}$ $V^2$ | $2.57x10^{-7}$±$6.17x10^{-8}$ $V^2$ |
| 6-12 | $2.93x10^{-7}$±$9.54x10^{-8}$ $V^2$ | $3.98x10^{-7}$±$9.76x10^{-8}$ $V^2$ |
| 15-30 | $9.47x10^{-8}$±$2.94x10^{-8}$ $V^2$ | $1.19x10^{-7}$±$2.68x10^{-8}$ $V^2$ |
| 40-70 | $5.43x10^{-8}$±$1.64x10^{-8}$ $V^2$ | $8.61x10^{-8}$±$1.41x10^{-8}$ $V^2$ |
| 80-120 | $1.31x10^{-8}$±$3.17x10^{-9}$ $V^2$ | $2.22x10^{-8}$±$3.60x10^{-9}$ $V^2$ |

| | | | | | |
|---|---|---|---|---|---|
| **B, inset** | Power Area Ratio<br>N=7-8 | unpaired 2-tail t-test | Ctrl vs DTA | p=0.638 | t=0.4822, df=13 |

| | Control | DTA |
|---|---|---|
| 2-5/6-12 Ratio | 0.75±0.10 | 0.68±0.08 |

| | | | | | |
|---|---|---|---|---|---|
| **D** | Ctrl Power Area Time<br>Ctrl N=5-8 | Mixed-effects analysis | baseline RM vs timebins | Frequency | **F(4,35)=4.733; p=0.0037** |
| | | | | Time | **F(2.179,62.65)=6.397; p=0.0023** |
| | | | | Frequency x Time | **F(16,115)=2.891; p=0.0005** |

| | 0-30 min | 30-60 min | 60-90 min | 90-120 min |
|---|---|---|---|---|
| 2-5 | 6.3±9.1% | 21±11.6% | 18.7±10.2% | 13.1±12.3% |
| 6-12 | -23.2±5.2% | 13.3±17% | 6.2±9.1% | 6.3±9.9% |
| 15-30 | 4.2±6.3% | 14.1±16.9% | 16.4±10.7% | 14.9±11.5% |
| 40-70 | 128.8±48.1% | 45.2±29.9% | 48.2±24.4% | 45.1±16.9% |
| 80-120 | 45.9±12.6% | 33.1±19.1% | 23.9±8.5% | 27.8±6.2% |

| | | | | | |
|---|---|---|---|---|---|
| **D, inset** | Ctrl Ratio time<br>Ctrl N=5-8 | Mixed-effects analysis | baseline RM vs timebins | Time | F(1.464,9.152)=2.859; p=0.117 |

| | Baseline | 0-30 min | 30-60 min | 60-90 min | 90-120 min |
|---|---|---|---|---|---|
| 2-5/6-12 | 1.4±0.2 | 1.6±0.1 | 1.6±0.1 | 1.5±0.1 | 1.4±0.1 |

| | | | | | |
|---|---|---|---|---|---|
| **F** | DTA Power Area Tim<br>DTA N=7 | 2way ANOVA | baseline RM vs timebins | **Frequency x Time** | **F(16,120)=5.793, p<0.0001** |
| | | | | Frequency | F(4,30)=2.536, p=0.0607 |
| | | | | Time | **F(2.355,70.65)=3.905, p=0.0191** |
| | | | | Mouse | **F(30,120)=6.491, p<0.0001** |

| | 0-30 min | 30-60 min | 60-90 min | 90-120 min |
|---|---|---|---|---|
| 2-5 | 1.6±14.5% | 62.5±19.7% | 40±12% | 43.9±12.3% |
| 6-12 | -19±9.2% | 25.4±17.4% | 19.7±15.6% | 34.6±12.3% |
| 15-30 | 0.2±8.8% | 7.7±7.3% | 0.5±4.7% | 8.9±4.3% |
| 40-70 | 18.1±12.9% | -11.9±13.3% | -10.4±12% | -12.5±12.5% |
| 80-120 | 13.4±10.5% | -5.2±11% | -7.6±9.1% | -10.6±10.6% |

| | | | | |
|---|---|---|---|---|
| DTA Ratio time<br>DTA N=7 | 1way ANOVA | baseline RM vs timebins | Time | F(1.539,9.233)=3.608, p=0.0775 |
| | | | Mouse | **F(6,24)=21.71, p<0.0001** |

| | Baseline | 0-30 min | 30-60 min | 60-90 min | 90-120 min |
|---|---|---|---|---|---|
| 2-5/6-12 | 1.5±0.2 | 1.8±0.2 | 2±0.2 | 1.9±0.3 | 1.6±0.2 |

| | | | |
|---|---|---|---|
| **not shown** | Ctrl vs DTA<br>N=5-8 | Multiple unpaired t-test | No significant discoveries |

| | | | | | | |
|---|---|---|---|---|---|---|
| **Figure 8** | **A** | **Open Field Duration** | 2 way ANOVA | Control vs DTA | **Interaction** | F(1,50)=0.082, p=0.776 |
| | | | | | Position | **F(1,50)=1243, p<0.0001** |
| | | | | | Treatment | F(1,50)=0.925, p=0.341 |

| | Control n=25 | DTA n=27 |
|---|---|---|
| Periphery | 514.0±7.5s | 510.6±9.3s |
| Center | 86.0±7.5s | 89.4±9.3s |

| | | | | | |
|---|---|---|---|---|---|
| **B** | **Light Dark Duration** | 2 way ANOVA | Control vs DTA | **Interaction** | F(1,50)=0.016, p=0.901 |
| | | | | Position | **F(1,50)=29.43, p<0.0001** |
| | | | | Treatment | F(1,50)=0.415, p=0.523 |

| | Control n=25 | DTA n=27 |
|---|---|---|

| | | |
|---|---|---|
| Light | 232.0±10.6s | 231.9±12.5s |
| Dark | 320.7±12.1s | 324.7±12.5s |

| | | | | | |
|---|---|---|---|---|---|
| C | **EPM Duration** | 2 way ANOVA | Control vs DTA | **Interaction** | F(2,100)=0.33, p=0.722 |
| | | | | Position | **F(1.43,71.67)=131.3, p<0.0001** |
| | | | | Treatment | F(1,50)=0.052, p=0.821 |

| | Control n=25 | DTA n=27 |
|---|---|---|
| Open | 119.8±15.3s | 121.6±14.3s |
| Closed | 294.7±18.5s | 287.1±19.2s |
| Intersection | 71.7±6.5s | 86.5±9.3s |

| | | | | | |
|---|---|---|---|---|---|
| D | **Social Interaction Du** | 2 way ANOVA | Control vs DTA | **Interaction** | F(2,100)=0.264, p=0.769 |
| | | | | Position | **F(1.28,64.17)=120.3, p<0.0001** |
| | | | | Treatment | F(1,50)=2.49, p=0.121 |

| | Control n=25 | DTA n=27 |
|---|---|---|
| Social Chamber | 277.0±13.2s | 269.5±9.8s |
| Center Chamber | 92.7±4.5s | 96.1±5.5s |
| Empty Chamber | 206.0±12.1s | 215.2±9.6s |

| | | | | | |
|---|---|---|---|---|---|
| E | **FC Freezing** | Mixed-effects analysis | Control vs DTA | Time | **F(5,166)=149.3, p<0.0001** |
| | | | | Treatment | **F(1,34)=18.29, p=0.0001** |
| | | | | **Interaction** | **F(5,166)=14.35, p<0.0001** |

| | Control n=18 | DTA n=16-18 |
|---|---|---|
| 30 s | 1.8±0.4% | 2.8±1.0% |
| 90 s | 3.1±0.6% | 2.4±0.8% |
| 150 s | 11.8±2.4% | 7.6±1.9% |
| 210 s | 32.0±3.7% | 18.2±2.7% |
| 270 s | 45.7±3.5% | 28.4±3.5% |
| 330 s | 58.9±3.3% | 30.5±3.9% |

| | | | | | |
|---|---|---|---|---|---|
| F | **FC Contextual** | Mixed-effects analysis | Control vs DTA | Time | **F(4,134)=8.59, p<0.0001** |
| | | | | Treatment | **F(1,34)=15.08, p=0.0005** |
| | | | | **Interaction** | F(4,134)=2.31, p=0.061 |

| | Control n=18 | DTA n=17-18 |
|---|---|---|
| 30 s | 26.9±6.7% | 14.8±3.4% |
| 90 s | 28.7±5.8% | 18.1±2.5% |
| 150 s | 39.8±4.7% | 26.2±2.4% |
| 210 s | 46.0±4.5% | 21.1±3.3% |
| 270 s | 50.4±5.2% | 24.3±3.5% |

| | | | | | |
|---|---|---|---|---|---|
| G | **FC Cued** | Mixed-effects analysis | Control vs DTA | Time | **F(4,136)=2.49, p=0.046** |
| | | | | Treatment | **F(1,34)=10.49, p=0.003** |
| | | | | **Interaction** | **F(4,136)=3.64, p=0.008** |

| | Control n=18 | DTA n=18 |
|---|---|---|
| 30 s | 25.0±4.0% | 19.0±2.7% |
| 90 s | 31.6±4.3% | 23.0±3.5% |
| 150 s | 37.7±4.5% | 22.6±2.8% |
| 210 s | 33.3±4.4% | 22.8±3.1% |
| 270 s | 42.0±5.7% | 16.5±1.9% |

| | | | | | | | |
|---|---|---|---|---|---|---|---|
| Figure S2 | B | Power Area | two-tailed t-test | | | | |
| | | | | uV^2 | Control n=13, N=4 | DTA n=17, N=4 | unpaired two-tail t-test | |
| | | | 3 - 6 Hz | | 0.026±0.011 | 0.04±0.013 | 0.4401 | t=0.7832, df=28 |
| | | | 15 - 80 Hz | | 0.034±0.006 | 0.074±0.016 | **0.0473** | t=2.075, df=28 |
